# Linking Polysaccharide Structure, Gelation Kinetics, and Function in Dynamic Acylhydrazone Hydrogels

**DOI:** 10.64898/2026.09.25.752777

**Authors:** Fereshteh Kazemi-Aghdam, Zuzana Varchulova, Jovana Zvicer, Jasmina Stojkovska, Aneta Křížová, Sahar Dinparvar, Lucy Vojtová, Luboš Danišovič, Igor Lacík, Abolfazl Heydari

## Abstract

Dynamic covalent hydrogels formed through reversible acylhydrazone crosslinking have emerged as promising injectable biomaterials. However, a fundamental gap remains in understanding how the macromolecular structure of oxidized polysaccharides (OxPs) governs gelation kinetics and how these kinetics pathways translate into material properties and cellular responses. We address this question by developing an acylhydrazone hydrogel library composed of alginate adipohydrazide crosslinked with oxidized alginate (OxA) or oxidized dextran (OxD), two reactive aldehyde-bearing polymers with comparable chemical functionality but fundamentally distinct backbone structure. By varying polysaccharide type, oxidation degree, and reaction pH, we decoupled the effects of chemical functionality from macromolecular structure and established quantitative structure–kinetics–property–function relationships. OxD-based hydrogels undergo rapid, largely pH-independent gelation, whereas OxA-based systems display pronounced pH-dependent kinetics with significantly delayed network formation under physiological pH. These differences in gelation kinetics and OxPs macromolecular structures lead to marked variations in hydrogel mechanics, including stiffness, stress relaxation, stability, injectability, and post-injection recovery. Importantly, differences in gelation kinetics modulate cell– matrix interactions in three-dimensional culture. Slowly forming OxA hydrogels maintained rounded chondrocyte shape, while rapidly gelling OxD networks induced transient cell elongation. Mesenchymal stem cells displayed similar shape regardless of gelation kinetics, indicating cell-type-specific responses to matrix formation dynamics.

## 1. Introduction

Dynamic covalent hydrogels represent a versatile class of polymeric biomaterials that combine the structural robustness of covalently crosslinked networks with the dynamic adaptability required for biological applications. These hydrogels consist of polymer networks crosslinked through reversible covalent bonds that can exchange dynamically, allowing network rearrangement while maintaining overall connectivity.^[1]^ Consequently, they exhibit time-dependent mechanical properties, including viscoelasticity, stress relaxation, and self-healing, closely resembling the dynamic behavior of the native extracellular matrix (ECM).^[2]^ Their biological performance is governed by the kinetics and thermodynamic stability of the reversible crosslinks, which regulate network reorganization and, in turn, determine macroscopic properties such as stiffness, stress relaxation, and structural recovery under mechanical loading.^[3]^ Importantly, this coupling between molecular dynamics and macroscopic mechanics plays a critical role in modulating cell–matrix interactions^[2a, 2b, 4]^ and provides a promising platform for tissue engineering^[5]^ and advanced in vitro models,^[4b, 5a, 5d]^ where mechanically adaptive and biomimetic microenvironments are required. Among the different dynamic covalent chemistries explored, hydrazone-based systems have attracted particular attention because they provide a favorable balance between bond stability and reversibility under aqueous conditions, making them especially attractive for biomedical hydrogel design.^[3a, 6]^

Hydrazone hydrogels are formed through condensation reactions between hydrazine or hydrazide-containing functional groups (R–NH–NH_2_) and carbonyl-containing compounds (R′–CHO), resulting in the formation of reversible hydrazone-type linkages (R–NH–N=CH–R′). These dynamic covalent bonds serve as the primary crosslinking points within the hydrogel network and largely determine its structural and mechanical behavior. The reaction mechanism involves nucleophilic attack of the terminal nitrogen on the electrophilic carbonyl carbon of the aldehyde, followed by elimination of water to generate the characteristic C=N bond. Under aqueous conditions, these reactions remain reversible, establishing a dynamic equilibrium between bond formation and dissociation. Consequently, the overall network behavior is governed by the balance between the forward (formation) reaction rate, the backward (dissociation) reaction rate, and the resulting equilibrium constant.^[3a]^ The reversible nature of hydrazone crosslinks allows the network to undergo molecular rearrangement after gel formation, thus, hydrazone hydrogels exhibit dynamic mechanical properties. These characteristics make them particularly well suited for minimally invasive cell delivery and other injectable biomaterial applications.^[6]^ Furthermore, the dynamic nature of hydrazone bonds and the continuous rearrangement of the network in response to cell-generated forces create conditions that modulate cell–matrix interactions and regulate cellular morphology and matrix remodeling within three-dimensional environments.^[7]^

The macroscopic properties of hydrazone hydrogels can be governed by the macromolecular architecture of hydrazone counterpart-bearing components, which strongly influence hydrogel formation, network organization, and viscoelastic behavior.^[3b, 8]^ The structure of the polymer network, including linear, branched, side-chain-functionalized, and star-shaped configurations, determines crosslinking density, chain mobility, and stress distribution throughout the hydrogel network. Polymer chains may be crosslinked through low-molecular-weight linear crosslinkers, multifunctional star-shaped molecules, or direct side-chain reactions between polymer backbones, yielding networks with markedly different mechanical characteristics. For example, hydrogels based on two-arm, four-arm, or eight-arm PEG derivatives exhibit distinct stiffness and relaxation profiles due to differences in network connectivity of polymer chains and effective crosslink density.^[5a, 6b, 8b]^ Side-chain architectures often produce more heterogeneous and dissipative networks with slower stress relaxation, whereas star-shaped systems typically generate mechanically stronger structures with enhanced elasticity and faster relaxation times.^[8a]^ In addition, hydrogel properties are further influenced by parameters such as stoichiometry between reacting amine and aldehyde functional groups,^[9]^ polymer concentration,^[5d, 10]^ and functional group valency.^[3b, 6a, 8a, 11]^ These structural factors collectively govern the balance between network mobility, highlighting the importance of macromolecular design in controlling the dynamic mechanical properties of hydrazone hydrogels.

At the molecular scale, the chemical structure of the aldehyde^[5a, 5b, 5d, 12]^ and amine-containing components (hydrazine and hydrazide-functionalized molecules)^[3a, 13]^ play a decisive role in regulating hydrazone bond dynamics and, consequently, the mechanical behavior of the resulting hydrogel. Variations in chemistry influence both the rate of bond exchange and the thermodynamic stability of the formed hydrazone-type linkages. In particular, aliphatic and aromatic aldehydes exhibit markedly different reactivities due to differences in electronic effect and resonance stabilization.^[5a]^ Aliphatic aldehydes generally form hydrazone bonds with more exchange dynamics (particularly the rate of dissociation), resulting in highly dynamic networks characterized by rapid stress relaxation and enhanced matrix adaptability, whereas aromatic aldehydes stabilize the C=N linkage through π-conjugation, thereby reducing exchange dynamics and producing mechanically more persistent hydrogels with prolonged relaxation behavior. These differences between aliphatic and aromatic aldehydes directly influence cellular responses, as the more dynamic aliphatic hydrazone networks better accommodate cell-generated forces, thereby promoting cell function and ECM deposition compared with the more static aromatic-based networks.^[5b, 5d]^

Among hydrazone-based dynamic covalent systems, acylhydrazones represent a particularly important subclass in which hydrazide-functionalized molecules (R–CO– NH–NH_2_) react with aldehydes (R′–CHO) to form reversible acylhydrazone bonds (R– CO–NH–N=CH–R′). They generally exhibit higher thermodynamic stability and less exchange dynamics than hydrazones because of conjugation of hydrazide with the adjacent carbonyl group.^[3a, 6]^ As a counterpart, oxidized polysaccharides (OxPs) represent a major class of reactive aliphatic aldehyde precursors for the formation of acylhydrazone-based hydrogels. OxPs are typically generated through periodate oxidation of polysaccharides, which introduces reactive aldehyde groups along the carbohydrate backbone. Common examples include oxidized alginate (OxA)^[6, 14]^ and oxidized dextran (OxD),^[15]^ both of which readily react with hydrazide-containing crosslinkers such as adipic acid dihydrazide (ADH)^[3a, 6, 9a, 10b, 14b]^ or PEG-dihydrazide ^[9a, 11]^ to form dynamic hydrogel networks. In some systems, multifunctional hydrazide-bearing polymers are also employed, enabling side-chain crosslinking architectures.^[10b]^ Although these hydrogels rely on the same underlying acylhydrazone chemistry, substantial differences in gelation behavior have been reported depending on the oxidized polysaccharide employed. Notably, OxA-based hydrogels generally exhibit significantly slower gelation at neutral pH than OxD-based systems, despite the presence of identical reactive aldehyde groups. Previous studies reported that OxA-ADH hydrogels require approximately 45 min to gel at pH 7.4,^[6a]^ whereas OxD-based acylhydrazone hydrogels can form within minutes even under near-neutral or basic conditions.^[15-16]^ These observations indicate that hydrogel formation is not solely dictated by bond-forming chemistry but is also strongly influenced by the structural characteristics of the polysaccharide backbone.

In this work, we address a fundamental and underexplored aspect of dynamic covalent hydrogel design: the extent to which OxPs backbone structures, independent of aldehyde functionality, governs acylhydrazone network formation and the resulting material properties. To this end, we developed a controlled hydrogel library based on Alg-ADH crosslinked with either OxA or OxD, two systems with comparable aldehyde reactivity but distinct macromolecular backbones, namely linear uronic acid-based chains and α-(1→6)-linked glucose structures, respectively. Although previous studies have reported differences in gelation behavior between OxA-and OxD-derived hydrogels, these findings remain largely descriptive and are often influenced by variations in experimental conditions, particularly the pronounced pH sensitivity of acylhydrazone chemistry. Here, we establish a systematic framework linking backbone architecture to molecular exchange dynamics, network formation, and viscoelastic properties under physiologically pH, demonstrating how molecular-level differences translate into macroscopic properties such as stiffness, stress relaxation, and structural stability. Finally, we evaluate the biological relevance of these hydrogels in three-dimensional cell culture, showing how backbone-dependent network dynamics influence the mesenchymal stem cells (MSCs) and chondrocytes response, including viability, spatial distribution, and maintenance of morphology following encapsulation and injection.

## 2. Experimental

### 2.1. Materials

The pharma-grade sodium alginate (NaAlg) Protanal® LF 10/60 with a G content of 60 mol% and weight-average molecular weight, *M*_w_, of 100 kg·mol^−1^, was obtained from DuPont. The reported G content was provided by the manufacturer. *M*_w_ values were determined by the size-exclusion chromatography (SEC) using the multi-angle laser light scattering and refractive index detectors.^[17]^ Dextran (Dex) from Leuconostoc spp, *M*_w_ ∼ 100–150 kg·mol^-1^ as provided by the manufacturer, was purchased from Sigma-Aldrich. The *M*_w_ of 81 kg·mol^−1^ was determined by the SEC using the multi-angle laser light scattering and refractive index detectors. Adipic dihydrazide, 4-(4,6-dimethoxy-1,3,5-triazin-2-yl)-4-methylmorpholinium chloride (DMTMM, ≥95.0%), methyl red (0.04% in water), and phenol red (0.04% in water) were purchased from TCI Europe N.V. Sodium(meta)periodate (NaIO_4_, ≥99.8%) was purchased from Sigma-Aldrich. Dialysis tubing cellulose membrane with a molecular weight cut-off of 14 kg·mol^-1^ was purchased from Sigma-Aldrich, and dialysis tubing with a molecular weight cut-off of 3.5 kg·mol^−1^ (Spectra/Por^®^) was purchased from Spectrum Laboratories Inc. Deuterium oxide (D_2_O, 99.90%) was purchased from Eurisotop. Gibco™ Dulbecco’s Modified Eagle Medium (DMEM, low glucose), Gibco™ Fetal Bovine Serum (FBS), Gibco™ Penicillin/Streptomycin (10.000 U·mL^-1^), Gibco™ phosphate-buffered saline (PBS), Gibco™ Trypsin-EDTA (0.05%, phenol red), and Invitrogen™ LIVE/DEAD™ viability/cytotoxicity kit for mammalian cells were purchased from Thermo Fisher Scientific. Chondrocyte Growth Medium (C-27101) was purchased from Sigma-Aldrich (Merck KGaA, Darmstadt, Germany).

### 2.2. Polymer synthesis and characterization

#### 2.2.1. Alginate adipohydrazide (Alg-ADH)

Alg-ADH was synthesized following a previously reported method for the amidation of NaAlg mediated by DMTMM.^[18]^ 2.4 g of NaAlg (12 mmol uronate units, 1 eq of COO^-^) was dissolved in 120 mL of water. Then, 10.45 g of adipic dihydrazide (60 mmol, 10 eq of NH_2_ groups) was added to the NaAlg solution. Subsequently, 1.65 g of DMTMM (6 mmol, 0.5 eq) was introduced in three equal portions over 3 h at 55 °C. The reaction mixture was then maintained under continuous magnetic stirring at 600 rpm and 55 °C for 24 hours. Purification of the product was carried out by dialysis using a cellulose membrane with a molecular weight cut-off of 14 kg·mol^-1^. Dialysis was performed first against 100 mmol·L^-1^ NaCl for 24 h, followed by water for an additional 48 h, with three changes per day. During dialysis, the pH of the dialysis solution was adjusted to 8 using NaOH solution. Alg-ADH was isolated by lyophilization. Alg-ADH samples were synthesized in five independent batches, summarized in Table S1, Supplementary Information.

#### 2.2.2. Oxidized alginate (OxA)

NaAlg was oxidized using a method described in the literature^[19]^ with slight modifications. 3 g of NaAlg (15 mmol, 1 eq of uronate units) was dissolved in 120 mL of water. To obtain different degrees of oxidation (DO), two separate reactions were carried out under identical conditions, differing only in the amount of NaIO_4_, which was dissolved in 30 mL of water and added in a single portion to the NaAlg solution. Specifically, 1.6 g of NaIO_4_ (7.5 mmol, 0.5 eq) was used to prepare oxidized alginate of high DO (OxA-H), while 0.384 g (1.5 mmol, 0.1 eq) was used to obtain oxidized alginate of low DO (OxA-L). The reaction proceeded at 600 rpm and ambient temperature for 6 h under continuous magnetic stirring. Purification of the reaction mixture was carried out by dialysis using a cellulose membrane with a molecular weight cut-off of 14 kg·mol^-1^. Dialysis was performed against water for 72 h, with three changes per day. During dialysis, the pH of the dialysis water was adjusted to 8 using NaOH solution. Finally, OxA was isolated by lyophilization. OxA samples were synthesized in five independent batches, summarized in Table S2, Supplementary Information.

#### 2.2.3. Oxidized dextran (OxD)

Dex was oxidized according to a previously reported method^[20]^ with slight modifications. 3 g of Dex (18.6 mmol, 1 eq of anhydroglucose units) was dissolved in 60 mL of water. To obtain different DO, two separate reactions were performed under identical conditions, varying only the amount of NaIO_4_, which was dissolved in 60 mL of water and added in a single portion to the Dex solution. Specifically, 1.98 g of NaIO_4_ (9.3 mmol, 0.5 eq) was used to prepare dextran of high DO (OxD-H), while 0.6 g (2.8 mmol, 0.15 eq) was used to obtain dextran of low DO (OxD-L). The reaction proceeded at 600 rpm and ambient temperature for 6 h under continuous magnetic stirring. Purification of the reaction mixture was performed by dialysis using dialyzing tubing with a molecular weight cut-off of 3.5 kg·mol^-1^ for OxD-L and 14 kg·mol^-1^ for OxD-H, respectively, with three changes per day. Finally, OxD was isolated by lyophilization. OxD samples were synthesized in five independent batches, summarized in Table S3, Supplementary Information.

#### 2.2.4. Characterization of polymers

The ^1^H NMR spectra were recorded using a 400 MHz spectrometer at 65 °C, without water signal suppression. For the measurements, the polymers were dissolved in D_2_O at a concentration of 10 mg·mL^-1^. The DS of Alg-ADH was determined by ^1^H NMR (*DS*_NMR_) and CHN elemental analysis (*DS*_EA_), and expressed as a percentage, following a previously reported method for modified NaAlg.^[18]^ A DS value of 14% indicates that 14% of the repeating units of NaAlg are functionalized with adipic dihydrazide, with a theoretical maximum DS of 100%. The *DS*_NMR_ values were calculated from the ratio of the integrated proton signals of the ADH side chains and the NaAlg backbone according to Equation 1:

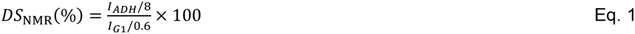

where *I*_ADH_ and *I*_G1_ correspond to the integrals of the methylene proton signals of the ADH substituent and the anomeric proton of the guluronic acid units in NaAlg, respectively, as previously described.^[18]^ The values of 8 and 0.6 represent the number of corresponding protons of ADH substituent and the fraction of anomeric protons associated with guluronic acid units in NaAlg, respectively.

The presence of nitrogen atoms in the synthesized Alg-ADH enabled the determination of the *DS*_EA_ by elemental analysis from the C/N atomic ratio. The *DS*_EA_ values were calculated according to Equation 2, which was derived from a polynomial fit correlating DS (in %) with the theoretical N/C atomic ratio (in %) (Figure S1, Supplementary Information).

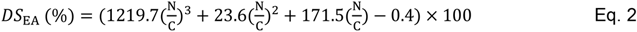

where *N*/*C* is the mole fraction ratio of nitrogen to carbon, measured by Equation 3:

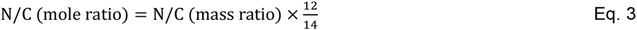

where 12 is the atomic mass of carbon, and 14 is the atomic mass of nitrogen.

##### DO of oxidized polymers

Schiff reagent assay was performed to obtain DO according to previously reported procedures.^[3a, 21]^ Briefly, Schiff reagent was reacted with aqueous solutions of OxPs for 1 h at room temperature. Then, UV–vis spectra were recorded to quantify aldehyde groups at the reagent’s characteristic wavelength of 550 nm. The calibration curve of OxD-H was obtained from the hydroxylamine hydrochloride assay (Table S4, Method S1, Supplementary Information). This calibration curve was subsequently used to determine the OD of the other samples (OxD-L, OxA-H, and OxA-L) based on their measured absorbance. DO of OxPs was calculated and is expressed as a percentage of the total repeating units in the polymer that were oxidized. The theoretical maximum DO is 100%, corresponding to oxidation of all repeating units, where each oxidized repeating unit generates two aldehyde functional groups.

##### Recovery yield of OxPs

Recovery yield of modified polymers, expressed as a percentage, was calculated using Equation 4, reflecting overall mass recovery after synthesis and purification.

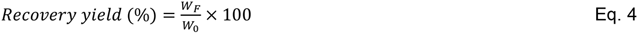

where *w*_F_ and *w*_0_ represent the weights of lyophilized product and the initial starting polymer, respectively. The oxidation does not significantly alter the molecular weight of repeating units for parent and oxidized polysaccharides, allowing direct comparison based on weight recovery.

##### Moisture content

Water content analysis was carried out using a moisture analyzer (VWR International, USA) based on thermogravimetric measurement. Samples were weighed and heated for 1 h at 105 °C in the instrument chamber. During the drying process, the decrease in sample weight was recorded until a stable weight was reached. The percentage of water content was calculated from the relative mass loss after drying. Measurements were conducted in triplicate.

##### Size Exclusion Chromatography (SEC)

SEC was employed to determine the molecular weights of NaAlg, Dex, OxD-H, OxD-L, OxA-H, and OxA-L. Measurements were performed using an aqueous SEC system equipped with PSS Suprema 10 μm guard and three columns (8 x 300 mm) with pore sizes 100, 1000 and 3000 Å, positioned in a Waters column heater module set to 40 °C, and an eluent consisting of 0.1 mol·L^-1^ Na_2_HPO_4_ and 200 ppm NaN_3_. Ethylene glycol was added to each sample as a flow marker to monitor the flow rate, which was set to 1 mL·min^-1^. Samples were dissolved in the SEC eluent, filtered through a 0.45 µm glass fiber syringe filter (Q-Max RR), and injected onto the columns using a 100 µL injection loop. Each sample was injected in duplicate. For NaAlg, OxA-H, and OxA-L, the absolute molecular weights were determined using a multi-angle laser light scattering detector (PSS SLD 7000, Polymer Standards Service) coupled with a differential refractive index detector (Waters 2414 DRI, Waters Corporation). Data analysis was performed using refractive index increment values of d*n*/d*c* = 0.136 mL·g^-1^ for pullulan (*M*_w_ = 110 kg·mol^-1^), which was used for detector calibration, d*n*/d*c* = 0.145 mL·g^-1^ for NaAlg, and d*n*/d*c* = 0.110 mL·g^-1^ for OxA-H, and OxA-L. For Dex, OxD-H, and OxD-L, the molecular weights were determined by calibration against narrow-distributed pullulan (Polymer Standards Service) and dextran (Biotika, a.s.) standards. WinGPC Unichrome software (Polymer Standards Service) was used for the determination of d*n*/d*c* values and for data acquisition and evaluation.

### 2.3. Gelation and hydrogel formation

#### 2.3.1. Polymer solutions

Polymer solutions were prepared at a total polymer concentration of 1.4 wt% in 0.9 wt% NaCl. The pH was adjusted to either 4.5 or 7.4 using 1 mol·L^-1^ HCl or 1 mol·L^-1^ NaOH (added volumes of HCl and NaOH solutions were negligible relative to the total solution volume). The pH values were measured using pH meter (pH 50+, XS Instruments, Italy). Hydrogel formulations were prepared by mixing solutions of Alg-ADH with OxA or OxD. Samples were labeled based on OxPs variables in the format of XY-Z, where X denotes A or D for OxA or OxD, respectively, Y denotes high, H, or low, L, degree of oxidation, and Z denotes acidic, A (pH 4.5), or neutral, N (pH 7.4), conditions of the OxPs solution.

#### 2.3.2. Gelation kinetics

Gelation kinetics was evaluated using both qualitative and quantitative approaches. Qualitative gelation time was determined by the inverted tube test, while quantitative analysis was performed using oscillatory time sweep rheology. Gelation was evaluated by mixing solutions of Alg-ADH with OxPs (OxA or OxD), at an amine-to-aldehyde molar ratio of [NH_2_] : [C=O] of 1 : 2 (Table 1). As a representative example, the AH-A sample (Table 1) was prepared by mixing 200 µL of Alg-ADH solution (2.8 mg of Alg-ADH, 14.5 µmol Alg-ADH, 1 eq of NH^2^, pH 7.4) with 73 µL of OxA-H solution (1.0 mg of OxA-H, 5.3 µmol OxA-H, 2 eq of C=O, pH 4.5), in 1.5 mL microcentrifuge tubes. The solutions were combined by vortexing (Vortex type) for 10 s under ambient conditions.

**Table 1.**
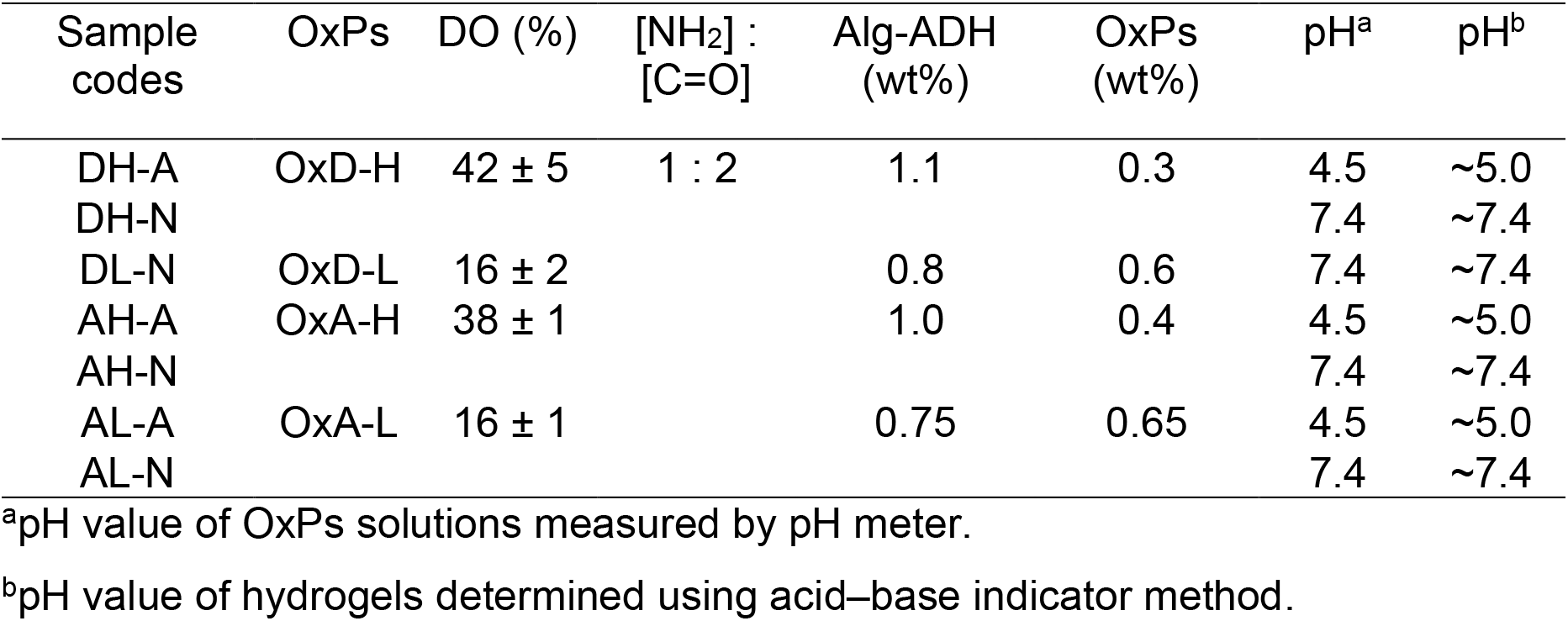
Formulations of acylhydrazone systems. Alg-ADH was used as the hydrazide-containing component in all systems. Polymer solutions were prepared at 1.4 wt% in 0.9 wt% NaCl. The total polymer concentration in hydrogel was 1.4 wt%. DS of Alg-ADH was 14 ± 2 mol%, and the pH of Alg-ADH solution was 7.4.

For qualitative assessment, the mixed solutions were transferred into glass vials (11.6 × 32 mm, 1.5 mL, flat bottom, clear; MACHEREY-NAGEL GmbH & Co. KG, Düren, Germany) and gently inverted at defined time intervals. Gelation time was defined as the time point at which no observable flow occurred upon inversion. Measurements were performed immediately after mixing (<30 s) and then at 5 min intervals. For rheological measurements, 200 µL of freshly mixed sample was loaded onto an Anton Paar MCR 302e rheometer equipped with a cone–plate geometry (20 mm diameter, 1° cone angle, 0.047 mm gap) and a Peltier temperature controller. Time-sweep measurements were initiated approximately 2 min after mixing (considered as dead time) and conducted at 25 °C for 120 min under oscillatory shear strain of 0.2% and 10 rad·s^-1^. Gel formation was monitored by tracking the evolution of storage modules (G′) and loss modules (G″) over time.

#### 2.3.3. Hydrogel preparation and processing

Hydrogels were prepared by mixing the polymer solutions using a dual-syringe mixing method based on the syringe-to-syringe transfer of polymer solutions described by Espona-Noguera et al.^[22]^ with slight modifications. Alg-ADH and OxPs solutions were loaded into separate polypropylene syringes and connected via a Luer-lock syringe connector (F/F adaptor, CELLINK AB, Gothenburg, Sweden). The solutions were mixed by three consecutive syringe-to-syringe transfers at a constant manual extrusion rate (total mixing time ∼5 s), followed by vortexing at 2500 rpm for 10 s under ambient conditions to prepare hydrogel precursors. For a representative formulation (AH-A), 1000 µL of Alg-ADH solution (14 mg of Alg-ADH, 9 µmol NH_2_, 1 eq; pH 7.4) and 325 µL of OxA-H solution (4.5 mg of OxA-H, 18 µmol C=O, 2 eq; pH 4.5) were transferred into separate 2 mL syringes and mixed.

Immediately after mixing (<30 s), hydrogel precursors were allocated to different experimental conditions. For rheological, uniaxial compression, and dynamic compression testing, hydrogels were loaded into a 1 mL insulin syringe mold, as described by Jin et al.^[23]^ For the injectability test, hydrogels were transferred into 1 mL insulin syringes. The molds or syringes were sealed to prevent dehydration of hydrogels and stored at ambient conditions for 24 h prior to testing.

#### 2.3.4. Sterile hydrogel preparation for testing under culture conditions

Polymer solutions were sterilized by filtration through 0.22 µm PES filters (TPP Techno Plastic Products AG, Trasadingen, Switzerland). Hydrogels were formed according to the method described in Section 2.3.3, with all procedures carried out under sterile conditions in a laminar-flow biosafety cabinet. Then, 200 µL of the hydrogel precursor mixture was dispensed into sterile 48-well plates. Hydrogels were overlaid with DMEM supplemented with 10% v/v FBS and 1% v/v penicillin/streptomycin (complete DMEM) and incubated in a standard cell culture incubator (Memmert GmbH, Germany) at 37°C in a humidified atmosphere containing 5% CO2. The culture medium was added approximately 5 min after mixing, except for AH-N hydrogels, which were allowed to crosslink for 50 min prior to medium addition. The samples were subsequently incubated for 28 days and used for compression testing.

#### 2.3.5. pH of hydrogels

The pH of the resulting hydrogels (Table 1) was qualitatively evaluated using an acid– base indicator method.^[24]^ Briefly, either methyl red or phenol red was added separately to the Alg-ADH solution at 1% (v/v) prior to gelation. Subsequently, the OxPs solution was introduced to initiate hydrogel formation. The pH of the formed hydrogels was then estimated from the color change of the indicators after gelation. Methyl red exhibits a color transition from red to yellow over the pH range of 4.2–6.2, whereas phenol red transitions from red (pH = 8.4) to yellow for pH < 6.8.

### 2.4. Oscillation rheology

Oscillatory rheology was performed using an Anton Paar MCR 302-e rheometer equipped with a Peltier temperature controller. A cone–plate geometry (20 mm diameter, 1° cone angle, 0.047 mm gap) was used for all measurements. For each experiment, 200 µL of hydrogel precursor was loaded onto the rheometer geometry, and the exposed sample edge was covered with silicone oil to prevent evaporation. Prior to testing, a 5 min time sweep was conducted at 0.2% strain and 10 rad·s^-1^ to confirm the equilibrium state, defined by constant viscoelastic moduli over time. Amplitude sweeps were performed over a strain range of 0.1-500% at 10 rad·s^-1^. The linear viscoelastic region (LVR) was defined as the strain range over which G′ remained within ± 5% of its plateau value. The yield point corresponded to the shear stress at the limit of the LVR, and the flow point was defined as the strain at which G″ exceeded G′. Frequency sweeps were conducted over an angular frequency range of 0.1-100 rad·s^-1^ at 0.2% strain. The loss factor (tan δ) was calculated as G″/G′. G′ recovery was evaluated using progressive strain–amplitude^[25]^ and cyclic step-strain^[26]^ recovery tests. In the progressive strain recovery tests, amplitude sweeps (0.1-500% at 10 rad·s^-1^) were followed by time sweeps at 10 rad·s^-1^ and 0.2% strain until the recovery of the initial G′. In the cyclic strain recovery tests, time sweeps were performed at 10 rad·s^-1^ by alternating between 0.2% strain (120 s) and 500% strain (60 s) for 10 cycles. Stress relaxation tests were performed using the method described in the literature,^[27]^ applying stepwise strains of 5, 10, 20, 30, 40, 50, 60, 70, 80, 90, and 100%, each maintained for 300 s. Steady shear viscosity was measured over a shear rate range of 0.1-100 s^-1^. All measurements were conducted at 37 °C. The stress relaxation response is described by Equation 5:^[4b, 5d]^

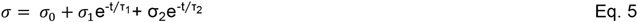

where σ_0_ is the non-relaxing plateau stress, τ_1_ and τ_2_, correspond to fast and slow relaxation timescales, and σ_1_ and σ_2_ denote their relative stress contributions.

### 2.5. Compression testing

Unconfined uniaxial compression of the hydrogels was performed using a TA.XTplus C Texture Analyser (Stable Micro Systems Ltd., Godalming, United Kingdom) equipped with a flat-ended cylindrical probe. Hydrogel specimens were prepared either by (i) sectioning hydrogels molded in 1 mL insulin syringes with 4.7 mm internal diameter or (ii) punching hydrogels formed and stored in well plates using a biopsy punch with a plunger of 4 mm internal diameter (Miltex®, Ted Pella Inc., Redding, CA, USA). The height of each specimen was 2.0 ± 0.5 mm as measured using a micrometer. Samples were placed on fine sandpaper (grit 320) to prevent slippage during compression testing, and 2–3 drops of water were applied around the specimen edge to minimize dehydration during testing. The initial height (*H*_0_) was defined as the displacement corresponding to a compressive force of 0.5 g and was automatically determined by the device. All tests were conducted under ambient conditions at a deformation rate of 1 cm·min^-1^, allowing accurate determination of equilibrium mechanical properties.^[28]^ Three different types of measurements were performed as previously reported:^[9d]^ (i) Compressive failure tests were performed by compressing the hydrogels up to 98% strain relative to *H*_0_; (ii) Cyclic loading–unloading tests consisted of 10 consecutive cycles at a maximum compressive strain of 20% relative to *H*_0_, the compressive modulus was calculated from the slope of the stress–strain curve within the low-strain regime (<10% strain relative to *H*_0_); (iii) Stress-relaxation tests were conducted by applying a constant compressive strain of 15% and recording stress decay for 300 s. The stress relaxation time was defined as the time required for the stress to decrease to 50% of its initial value, as described in the literature.^[2b]^

### 2.6. Dynamic compression testing in a bioreactor with dynamic compression and medium perfusion

Mechanical evaluation under in vivo*-*like condition were performed on AH-N and DH-N hydrogels using a custom bioreactor system enabling simultaneous uniaxial mechanical loading and medium perfusion.^[29]^ Hydrogel specimens (12 mm diameter, 3 mm height) were placed in bioreactor chambers (16 mm inner diameter, 6 mm inner height) and incubated for 7 days at 37 °C and 5% CO_2_. Three chambers were operating in parallel, each connected to an independent recirculation loop containing 10 mL of complete DMEM supplemented with 50 µg·mL^-1^ ascorbic acid. The culture medium was continuously perfused at a flowrate of 0.7 mL·min^-1^, corresponding to a superficial velocity of 150 µm·s^-1^ using a multichannel peristaltic pump (Masterflex, Cole-Parmer, IL). Dynamic compression was applied for 1 h/ day at a loading rate of 337.5 μm·s^-1^, frequency 0.56 Hz, with total displacement of 0.3 mm, corresponding to approximately 10% strain.

The stress values were calculated using the initial plunger cross-sectional area. The compressive modulus values were calculated as described in Section 2.5. Control samples were cultured under static conditions in 6-well plates in 10 mL complete DMEM supplemented with 50 µg·mL^-1^ ascorbic acid each at 37 °C and 5% CO2. Each experiment was performed using freshly prepared hydrogels obtained from a single polymer batch, with three independent replicates.

### 2.7. Injectability

The injectability of the acylhydrazone hydrogels was evaluated using a syringe extrusion test performed on a TA.XTplus C Texture Analyser (Stable Micro Systems Ltd., Godalming, UK), as described previously.^[30]^ A 1 mL insulin syringe filled with hydrogel and fitted with a 21 G needle was positioned beneath the probe. The syringe was fixed using side clamps, while the plunger was allowed to move freely. The probe was moving at a constant rate of 1 mm·s^-1^ to displace the syringe plunger. The extrusion force was recorded as a function of plunger displacement. A syringe filled with distilled water was used as a control under identical conditions. All measurements were performed in triplicate.

To evaluate the recovery of viscoelastic properties after injection, hydrogels were extruded through a 21 G needle using the syringe extrusion setup described above. The injected hydrogels were collected and allowed to recover for 30 min at ambient conditions in closed sample vials. Subsequently, oscillatory frequency sweep measurements were performed as described in Section 2.4. Recovery of viscoelastic properties after injection was quantified by calculating the ratio of post-injection to pre-injection G’ (G′_post_-/G′_pre-injection_) at 10 rad·s^-1^.

### 2.8. Printability

Hydrogel inks were prepared by mixing solutions of Alg-ADH with OxPs at an amine-to-aldehyde molar ratios of [NH_2_] : [C=O] of 1 : 2 (Table 1), 1 : 1, and 2 : 1 (Table 2) according to the method described in Section 2.3.3. The hydrogel inks were loaded immediately after mixing into 3 mL syringes fitted with a 22 G nozzle. Approximately 10 min after mixing the polymer solutions, the printing was performed under ambient conditions via pneumatic extrusion using an extrusion-based Cellink BioX bioprinter. For the sample AH-N-R2, the printing was initiated 20 min after mixing due to the slower gelation kinetics. The printability was defined as the ability of the hydrogel ink to flow and form continuous filaments. Filament fusion and shape fidelity were qualitatively evaluated by printing square-lattice constructs with square pore geometry at 30% infill, consisting of three stacked layers forming 2 cm square structures. This qualitative assessment was based on the ability of the printed constructs to maintain the open spaces between filaments and preserve the square pore geometry without excessive spreading or filament merging.^[31]^ Representative printed constructs were documented and evaluated by digital photography.

**Table 2.** Formulations of acylhydrazone hydrogel inks. The polymer solutions were prepared at 1.4 wt% in 0.9 wt% NaCl. DS of Alg-ADH was 14 ± 2%, DO of OxD-H was 38 ± 1%, and DO of OxA-H was 42 ± 5%. pH of Alg-ADH solution was 7.4, and total concentration of polymers was 1.4 wt%.

| Sample code | OxPs | [NH<sub>2</sub>] : [C=O] | Alg-ADH (wt%) | OxPs (wt%) | pH<sup>a</sup> | pH<sup>b</sup> |
| --- | --- | --- | --- | --- | --- | --- |
| DH-N-R1 | OxD-H | 1 : 1 | 1.2 | 0.2 | 7.4 | ~7.4 |
| AH-A-R1 | OxA-H |  | 1.2 | 0.2 | 4.5 | ~5.0 |
| DH-N-R2 | OxD-H | 2 : 1 | 1.3 | 0.1 | 7.4 | ~7.4 |
| AH-A-R2 | OxA-H |  | 1.3 | 0.1 | 4.5 | ~5.0 |

<sup>a</sup>pH value of OxPs solutions measured by pH meter.
<sup>b</sup>pH value of hydrogels determined using acid–base indicator method.

### 2.9. Cell-based experiments

#### 2.9.1. Cell culture and expansion

Human adipose-derived mesenchymal stem cells (MCSs) (catalog number SCC038, Merck Millipore, Darmstadt, Germany) were used in this study. The cryopreserved cells (received at passage 2) were thawed and maintained as an adherent monolayer in complete DMEM. All sourced from catalog number SCC038, Merck Millipore (Darmstadt, Germany). The cultures were incubated under standard conditions at 37°C in a humidified atmosphere containing 5% CO_2_.

The cell culture medium was replenished every 2 days. Upon reaching approximately 80–90% confluence, the cell monolayer was rinsed with PBS and detached using Trypsin-EDTA. The enzymatic reaction was neutralized with complete DMEM, and the cell suspension was centrifuged at 230 × g for 10 min. The resulting cell pellet was resuspended in fresh complete medium and subcultured. For all subsequent downstream assays, MSCs at passages 4 and 5 were used to ensure phenotypic consistency and stability.

Cryopreserved human chondrocytes (5 × 10^5^ cells; Sigma-Aldrich, catalog number C-12710) were cultured in chondrocyte growth medium according to the manufacturer’s instructions. Cells were seeded into T-75 flasks at an initial density of 5 × 10^5^ cells·mL^- 1^ and incubated at 37 °C in a humidified atmosphere with 5% CO_2_ until they reached approximately 80% confluency. Cells were subsequently washed with PBS, detached using Trypsin–EDTA, centrifuged, resuspended in fresh complete medium, and counted prior to further experimental procedures.

#### 2.9.2. Preparation of cell-laden hydrogels

Polymer solutions were sterilized by filtration through 0.22 µm PES filters. Cells were suspended in Alg-ADH solution by gentle pipetting and subsequently mixed with the corresponding OxPs solution, referred to as the cell-laden hydrogel precursor, according to the protocol described in Section 2.3.4. MSCs were laden at a cell density of 1 × 10^6^ cells·mL^-1^ in DH-N, DL-N, AH-A, AL-A, AH-N, and AL-N formulations. Chondrocytes were encapsulated at a cell density of 2 × 10^6^ cells·mL^-1^ in AH-N and DH-N selected as representative hydrogels. Cell densities are reported with respect to the hydrogel volume after preparation. Immediately after mixing, the cell-laden hydrogel precursor was processed according to the specific test described below. MSCs-laden hydrogels were incubated in complete DMEM, whereas chondrocytes-laden hydrogels were incubated in chondrocyte growth medium at 37 °C in a humidified atmosphere with 5% CO_2_ until further analysis.

#### 2.9.3. Cytocompatibility and cell distribution

Cell-laden hydrogel precursors (200 µL) were dispensed into 48-well plates. Hydrogels were covered with complete DMEM, except for the AH-N sample, which was allowed to crosslink for 50 min prior to medium addition. The samples were incubated for 28 days at 37 °C in a humidified atmosphere containing 5% CO_2_ and were analyzed on days 1, 7, 14, and 28. Cells were stained using the LIVE/DEAD™ Viability/Cytotoxicity Kit according to the manufacturer’s protocol. Briefly, the cell-laden hydrogels were washed three times in PBS and incubated in a LIVE/DEAD staining solution containing calcein-AM and ethidium homodimer-1 (1 µL·mL^-1^ each in PBS) for 15 min at 37 °C. Samples were fully submerged during staining and subsequently rinsed once with PBS prior to imaging. Three independent hydrogels were analyzed for each time point.

Fluorescence imaging was performed using a confocal laser scanning microscope (LSM 900, Zeiss, Jena, Germany) equipped with 10× and 40× objectives. The 10× objective was used to assess cell distribution and viability, while the 40× objective enabled detailed evaluation of cell shape. Calcein-AM (live cells) was excited at 488 nm (emission 515 nm) and ethidium homodimer-1 (dead cells) at 543 nm (emission 635 nm). The entire hydrogel construct was scanned in the x-, y-, and z-dimensions to confirm uniform cell distribution. Subsequently, at least three regions at different x-, y-, and z-locations were imaged, and representative images are shown. Z-stacks were acquired from the top view and reconstructed for qualitative analysis. For detailed axial imaging, stacks covered the first 200 µm with 5 µm z-steps, whereas volumetric imaging spanned 1.4 mm with 40 µm z-steps. The total hydrogel thickness was approximately 2 mm.

#### 2.9.4. Quantitative phase imaging of chondrocytes

Quantitative phase imaging (QPI) was performed using a multimodal holographic microscope (Q-PHASE, Telight a.s., Brno, Czech Republic) equipped with 10× and 40× objectives. Chondrocyte-laden AH-N and DH-N hydrogels were prepared by mixing the hydrogel precursors with a chondrocyte suspension (section 2.9.2) and immediately loading the mixtures into µ-Slide I Luer channel slides (0.8 mm channel height, 200 µL volume; Ibidi GmbH, Germany), followed by the addition of Chondrocyte Growth Medium to the channel reservoirs. For live-cell imaging, the slides were maintained in the microscope environmental chamber throughout the experiment, and quantitative phase images were acquired every 10 min for 18 h. For long-term evaluation, chondrocyte-laden hydrogels were prepared in glass-bottom culture dishes (FluoroDish, World Precision Instruments, Europe), cultured under standard conditions for up to 21 days, and imaged at the designated time points using the same QPI system.

#### 2.9.5. Chondrocyte shape analysis

Chondrocyte shape was quantified using the deformation index X/Y, calculated as the ratio of the minor cell axis X to the major cell axis Y. A deformation index close to 1 indicates a rounded cell, whereas values below 1 indicate increasing cell elongation. Cell dimensions were measured from microscopy images using ImageJ software, and the deformation index was calculated from five randomly selected cells for each hydrogel and time point. Results are presented as mean ± SD.

#### 2.9.6. Post-injection viability of MSCs

For post-injection viability assessment, cell-laden hydrogel precursors of DH-N and AH-N formulations were loaded into standard insulin syringes fitted with a 21 G needle. The AH-N precursor was incubated at 37 °C for 50 min before injection, whereas DH-N was injected immediately after encapsulation. Subsequently, 200 µL of the resulting hydrogels were injected into 48-well plates. Complete DMEM was then added, and the samples were incubated for 24 h. Cell viability was assessed using LIVE/DEAD staining as described earlier (Section 2.9.3). Three independent injected hydrogels were prepared from the same precursors for each condition.

#### 2.9.7. Stability of MSCs-laden hydrogels

The MSCs-laden hydrogel precursors, including DH-N, DL-N, AH-A, AH-N, AL-A, and AL-N, were transferred into insulin syringe molds (Section 2.3.3). The molded cylindrical hydrogels with a 4.7 mm diameter, were subsequently sectioned into hydrogel discs with a height of 2.0 ± 0.5 mm, placed in 24-well plates, and incubated for up to 90 days in a cell culture incubator. At predefined time points on day 1, 7, 14, 21, 30, 40, 60, and 90, culture medium was removed, and top-view digital photographs were acquired to document visible changes in hydrogel dimensions. The culture medium was replaced twice weekly throughout the incubation period. Samples were classified as “GONE” upon complete dissolution, defined as the absence of any remaining hydrogel structure. Three independent hydrogels were analyzed at each time point.

For mechanical testing, MSCs-laden AH-N (AH-N@MSCs) and DH-N (DH-N@MSCs) hydrogels were prepared as described in section 2.9.2 and incubated for up to 28 days in a cell culture incubator. At predefined time points, on days 1, 7, 14, and 28, hydrogels were removed from the culture medium and subjected to compressive failure testing, as described in Section 2.5.

### 2.10. Repeatability

Repeatability was assessed across the experimental workflow by evaluating independent synthesis batches, hydrogel preparation, and rheological characterization. G’ values at strain 0.2% and at frequency of 10 rad·s^-1^ were used as the repeatability parameter and are summarized in Table S5, Supplementary Information. For all other experiments, measurements were performed in triplicate using freshly prepared hydrogels from a single synthesis batch, ensuring consistent material properties within each experimental set. This approach enables reliable assessment of hydrogel behavior while minimizing variability arising from differences in synthesis.

### 2.11. Statistical analysis

Statistical analysis was performed using OriginPro 2025 (OriginLab, Northampton, MA, USA). Data are presented as mean ± standard deviation (SD). For bioreactor experiments involving repeated measurements of hydrogels over time (n = 3), repeated-measures analysis of variance (RM-ANOVA) was performed, followed by Bonferroni-adjusted post hoc comparisons. For experiments involving independent groups (n = 3), one-way ANOVA followed by Tukey’s multiple comparison test was applied. A p-value <0.05 was considered statistically significant.

## 3. Results and discussion

### 3.1. Polymer library of acylhydrazone hydrogels

A polymer library incorporating two counterparts acylhydrazone-forming functional groups was developed based on NaAlg and Dex (Figures 1a and 1b). NaAlg was functionalized with ADH to yield Alg-ADH via DMTMM-mediated amidation of carboxyl groups, following a protocol adapted from our previous work.^[18]^ This modification introduced pendant hydrazide (R–CO–NH–NH_2_) functionalities along the polymer backbone. Alg-ADH was synthesized with a DS of 14 ± 2%, providing a sufficient density of reactive groups for efficient network formation, as reported previously in the literature.^[8a, 10b]^ Table S1 (Supplementary Information) summarizes the characterization data of independent synthesis batches of Alg-ADH. Structural characterization by ^1^H NMR spectroscopy confirmed successful functionalization. In the ^1^H NMR spectra (Figure 1c), NaAlg exhibits characteristic signals between 4.0 and 5.8 ppm corresponding to sugar ring protons, with the anomeric proton of guluronic acid (G1) at ∼5.2 ppm used as an internal reference for DS calculation. Following functionalization, new resonances appear at 1.6–2.3 ppm and 2.6–2.7 ppm, assigned to the methylene protons of the ADH moiety, confirming chemical structure of Alg-ADH.

**Figure 1.**
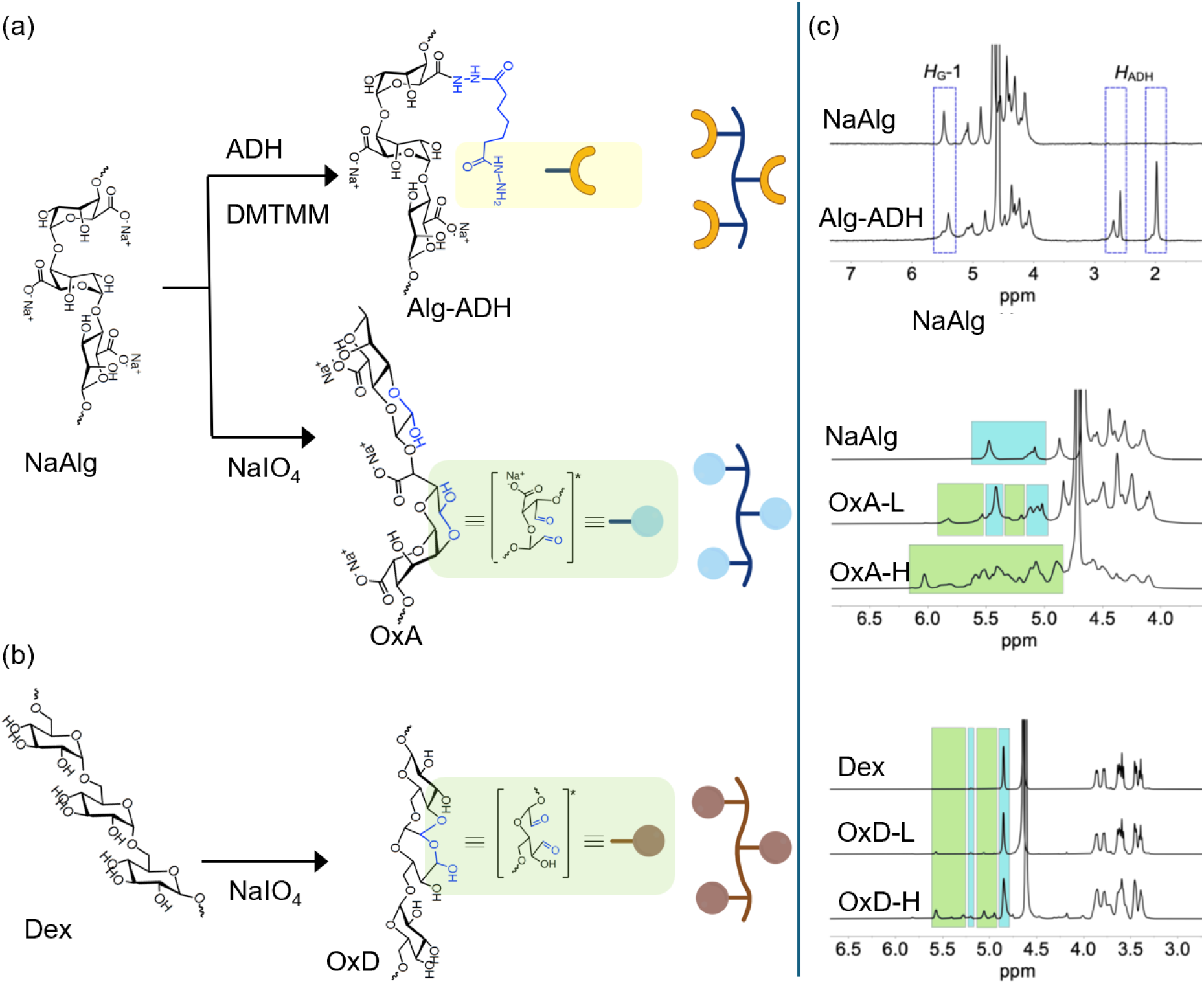
Schematic representation of (a) Alg-ADH and OxA synthesis from NaAlg, (b) OxD synthesis from Dex. The acylhydrazone counterpart in each polymer is indicated by a highlighted green square. Hemiacetal/hemialdal forms are referred to as “aldehyde groups” and are shown as reactive intermediates. (c) ^1^H NMR spectra (D2O, 600 MHz, 60 °C) of polymers. In Alg-ADH, the characteristic signals used for DS determination are labeled. In OxPs, the blue-highlighted region corresponds to protons of the main polymer backbone, and the green-highlighted region indicates newly formed protons after oxidation. No distinct assignment of signal is possible for OxA-H.

Aldehyde functionalities (R′–CHO) were introduced into NaAlg and Dex through periodate-mediated oxidation, producing OxA and OxD via cleavage of vicinal diols. In aqueous solution, these aldehyde groups exist predominantly in equilibrium with their intra- and intermolecular hemiacetal/hemialdal forms. As a result, the concentration of the free aldehyde species is typically too low for detection by ^1^H NMR, and the characteristic aldehyde proton resonance (∼9–10 ppm) is not observed, consistent with previous reports.^[20, 32]^ Despite this, these reactive moieties are referred to as “aldehyde groups” throughout this work for clarity and consistency in terminology. OxA and OxD were prepared with theoretical DO of 15 and 40%, based on the molar ratio of sodium periodate to the initial sugar units in Dex and NaAlg. These DO values enabled systematic variation in reactive group density while maintaining comparable crosslinking stoichiometry. These DO values were experimentally quantified using the Schiff reagent method, as summarized in Table 1 and Tables S4 and S6 (Supplementary Information). These DO values have previously been demonstrated to yield cytocompatible dynamic networks at an equimolar hydrazide-to-aldehyde ratio, exhibiting shear-thinning, self-healing, and stress-relaxation behavior for OxA hydrogels.^[6, 9a]^ However, these studies employed ADH as the hydrazide-containing crosslinker, rather than Alg-ADH.

The ^1^H NMR spectra of OxPs show characteristic changes compared to their native counterparts (Figure 1c). For NaAlg, signals are mainly observed between 4.4 and 5.5 ppm, including anomeric protons at ∼5.1 and ∼4.7 ppm, while Dex shows a dominant anomeric signal at ∼4.7 ppm and ring proton signals between 3.3 and 4.0 ppm. Upon oxidation, new signals attributed to hemiacetal proton environments, formed via reaction of aldehyde groups with neighboring hydroxyl groups, appear in both systems. Specifically, OxA shows additional broadened resonances in the 4.8–5.8 ppm region, whereas OxD displays several new signals in the ∼4.7–5.7 ppm range. These features significantly overlap with the polysaccharide backbone, making direct quantification of aldehyde groups by ^1^H NMR challenging, as widely reported.^[20, 32]^ Increasing the theoretical DO from 15 to 40% results in more pronounced peak broadening, reflecting increased backbone cleavage, reduced chain regularity, and enhanced structural heterogeneity. In OxA, the higher intensity and overlap of hemiacetal-associated signals at high DO hinder clear differentiation between oxidized and non-oxidized sugar units. In contrast, OxD retains a discernible anomeric signal, while the relative intensity of oxidation-related resonances increases with DO, enabling clearer qualitative assessment of the oxidation level. The experimentally determined DO values, measured using Schiff reagent assays, were 42 ± 5% for OxD-H, 16 ± 2% for OxD-L, 38 ± 1% for OxA-H, and 16% for OxA-L, which closely correspond to the theoretical DO values. Tables S2 and S3 summarize the characterization data for independent synthesis batches for OxA and OxD, respectively.

SEC analysis of molecular weights showed that increasing the DO reduced the molecular weight of OxPs. The Mw of OxPs were 8,500 ± 700 g·mol^-1^ for OxD-H, 30,000 ± 5,000 g·mol^-1^ for OxD-L, 19,400 g·mol^-1^ for OxA-H, and 27,000 g·mol^-1^ for OxA-L. This reduction in molecular weight upon oxidation is significant, nevertheless, resulting OxPs are still the long-chain polymers that are an important factor in enhancing the stability of imine-based dynamic bonds.^[33]^

### 3.2. Gelation kinetics in formation of acylhydrazone networks

The gelation kinetics of acylhydrazone-mediated networks, shown in Figure 2, was investigated to evaluate the influence of OxPs structure and crosslinking chemistry on network formation. Acylhydrazone network formation proceeds through reversible condensation between hydrazide and aldehyde functionalities (Figure 2a), a reaction known to be sensitive to both the chemical structure of the reactants and solution pH.^[3a, 3c, 34]^ These networks were formed by crosslinking Alg-ADH, which provides hydrazide functionalities along the polymer backbone, with either OxA or OxD as aldehyde-bearing counterparts (Figure 2b), enabling a systematic comparison of these two OxPs.

**Figure 2.**
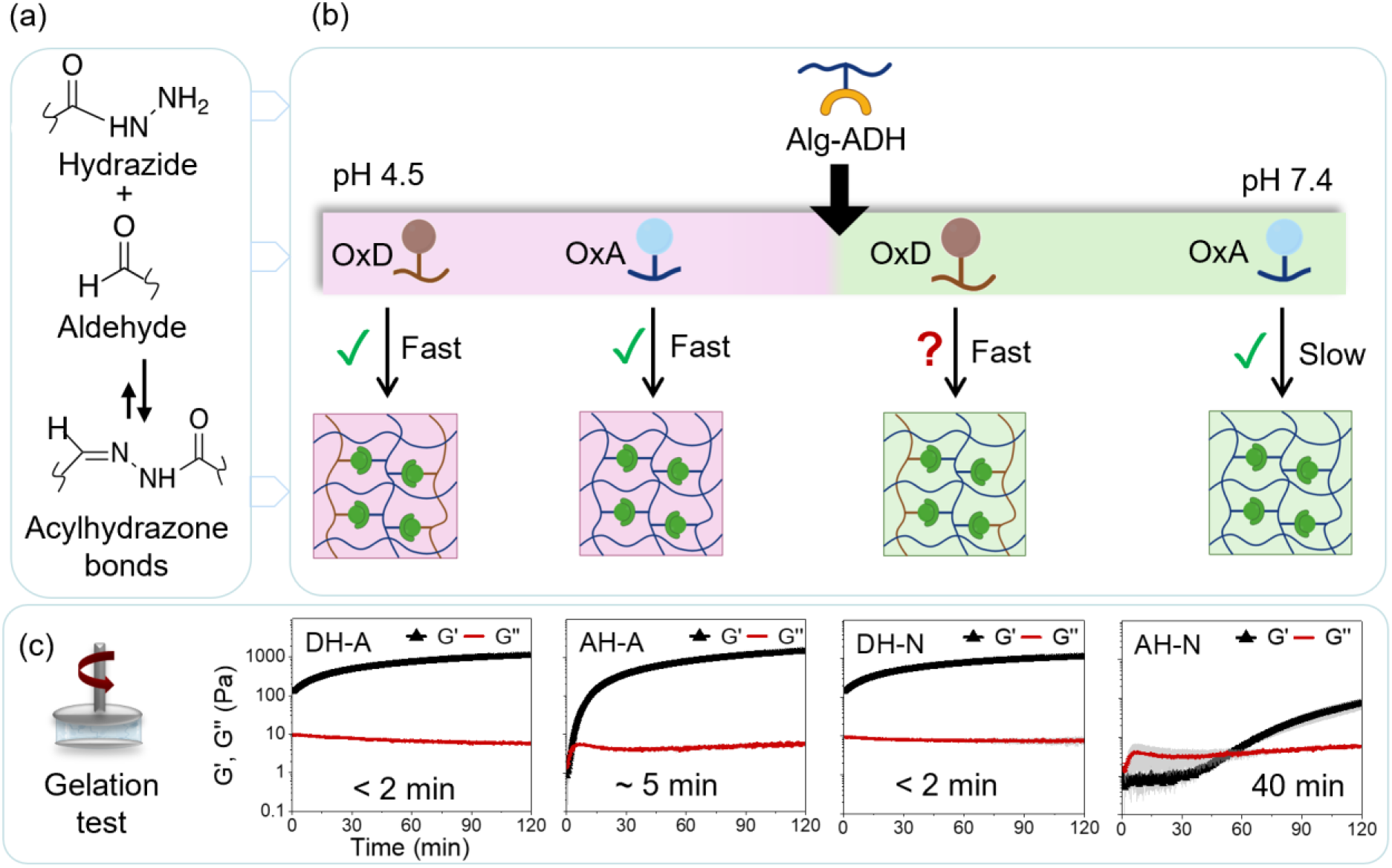
(a) Schematic illustration of reversible acylhydrazone bonds formation via the reaction between hydrazide and aldehyde functional groups. (b) Acylhydrazone network formation as a function of pH and the type of aldehyde-containing counterpart (OxA and OxD). (c) Representative oscillatory time-sweep measurements of the corresponding formulations performed at 0.2% strain and a frequency of 10 rad·s^-1^. Rheological measurements were performed in triplicate, with variability shown as shaded regions around the mean curves. Where shading is absent, the standard deviation was negligible.

Initial screening by vial inversion (Figure S2, Supplementary Information), followed by oscillatory time-sweep rheology, revealed pronounced differences in gelation behavior depending on both the OxPs and pH. OxA-based systems exhibited strongly pH-dependent gelation kinetics: under acidic conditions, gelation occurred within minutes (less than 5 min), whereas at neutral pH, network formation was significantly slower, requiring nearly one hour to form a gel. In contrast, OxD-based formulations with both high and low DO gelled rapidly, within seconds to only a few minutes (less than 5 min), under both neutral and acidic conditions. Rheological analysis (Figure 2c and Figure S3, Supplementary Information) confirmed these trends, where gelation was defined by the crossover of storage (G′) and loss (G″) moduli. OxD systems reach a solid-like response almost immediately within the experimental dead time (<2 min), while OxA systems showed a fast gelation under acidic conditions, and delayed network formation under neutral conditions.

Overall, gelation times across all formulations ranging from seconds to nearly one hour (Figure 2c and Figure S3) highlight the strong dependence of acylhydrazone network formation on both pH and OxPs backbone chemistry. The acceleration under acidic conditions is consistent with the proton-assisted mechanism of acylhydrazone bond formation, which proceeds faster at mildly acidic pH (∼ 4.5).^[34]^ At neutral pH, reduced proton availability slows the dehydration of the hemiaminal intermediate,^[35]^ making gelation more sensitive to polymer-dependent structural factors for OxA and OxD. The principal difference in molecular architectures of these two OxPs is that OxA is derived from a linear copolymer of uronic acids, whereas OxD is formed by oxidation of α-(1→6)-linked glucose units. This effect is pronounced for OxA-based systems (AH-A vs AH-N, Figure 2c), in agreement with previous reports describing slower gelation of OxA crosslinked using ADH at physiological pH (approximately 45 min to gel at pH 7.4).^[6a]^ Similarly, OxA-based hydrogels crosslinked via PEG-3,3′-dithiobis(propionohydrazide) have shown gelation times ranging from seconds to hours as pH increases from acidic to neutral values.^[11]^ Interestingly, OxD-crosslinked hydrogels (DH-A vs DH-N, Figure 2c) exhibited rapid gelation even under neutral conditions, suggesting that factors beyond proton availability contribute to their accelerated network formation. Consistent with this interpretation, previous studies have reported rapid gelation (<3 min) of acylhydrazone OxD hydrogels crosslinked with ADH in phosphate buffer at near-neutral or basic pH (pH 8.0).^[15a, 16]^

In addition, the influence of the DO on gelation kinetics was investigated. Reducing the DO altered both the relative proportions of Alg-ADH and OxD in the formulations, while maintaining constant overall polymer concentration and molar ratio of [NH_2_] : [C=O] of 1 : 2. As shown in Figure S3 (Supplementary Information), hydrogels prepared with a lower DO, e.g., DL-N and AL-A, exhibited rapid gelation and readily formed crosslinked networks, comparable to formulations prepared at higher DO (DH-N and AH-A) (Figure 2c). AL-N also exhibits a slow formation rate, comparable to that of AH-N prepared at higher DO. These observations indicate that observed gelation kinetics are independent of DO and are primarily governed by the chemical structure of the OxPs backbone.

### 3.3. Acylhydrazone hydrogel library and characterization states

The distinct gelation kinetics served as the primary design parameter for the systematic development and evaluation of an acylhydrazone hydrogel library, establishing correlations between crosslinking kinetics and bulk material properties (Figure 3). The hydrogels were formed using a dual-syringe mixing method in which Alg-ADH, serving as the hydrazide-containing component, was combined with aldehyde-functionalized OxPs to form dynamic acylhydrazone networks (Figure 3a). For cell-based studies, the Alg-ADH solution was mixed with either MSCs or chondrocytes prior to gelation.

**Figure 3.**
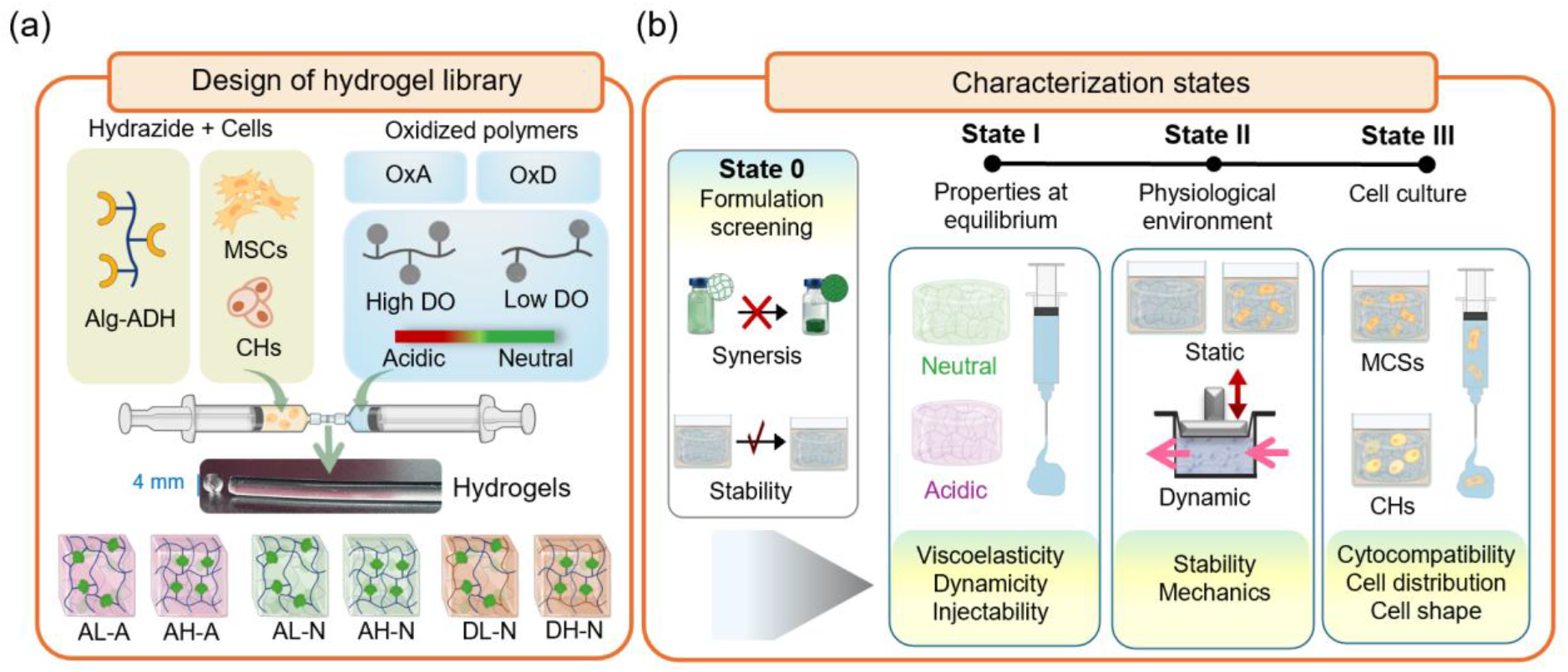
(a) Schematic overview of the design and (b) multi-stage characterization of the injectable acylhydrazone hydrogel library.

To form a structurally diverse hydrogel library, three key formulation parameters associated with the OxPs were varied (Figure 3a): (i) polysaccharide chemical structure (OxA vs OxD), (ii) crosslinking environment (acidic vs neutral conditions), and (iii) DO values (low vs high). This experimental design enabled a systematic investigation of how polymer structure and reaction conditions influence macroscopic hydrogel properties. As the hydrogel system is intended for cell culturing applications, formulations prepared under acidic conditions were not further considered, except for AH-A, which was retained as a representative sample because its gelation behavior differed markedly from that of AH-N. On the other hand, the effect of DO was specifically investigated in the OxD-based system by comparing DH-N and DL-N, particularly given the rapid gelation observed under neutral conditions for both formulations. The resulting hydrogel library comprised six formulations: AL-A, AH-A, AL-N, AH-N, DL-N, and DH-N (Figure 3a). Following gel formation, all formulations were subjected to an initial screening stage (State 0, Figure 3b), in which stability, syneresis behavior, and long-term storage stability were evaluated. Based on these criteria, four representative hydrogels (AH-A, AH-N, DL-N, and DH-N) were selected for further detailed characterization (State I-III, Figure 3b).

Given the known sensitivity of acylhydrazone bond formation to pH, the pH of the reaction environment during polymer solution mixing and gelation was verified using colorimetric pH indicators (Figure S4, Supplementary Information). Phenol red indicated that the Alg-ADH precursor solution was at neutral conditions (red to orange color, ∼pH 7.4) and retained this color after mixing with neutral OxA, confirming that gelation proceeded without measurable pH change. In contrast, mixing Alg-ADH with acidic OxA led to a clear shift to yellow color, consistent with mildly acidic conditions (pH < 6.8). This observation was further detailed by using the methyl red, which transitioned from yellow in the Alg-ADH solution to red upon addition of OxA-A, corresponding to a pH of approximately 4.5–5. These results confirm that the intended reaction environments are preserved during hydrogel formation, with AH-N systems remaining neutral and AH-A systems forming under acidic conditions.

The selected hydrogels were evaluated under conditions representing the network equilibrium state (State I, Figure 3b) and physiological environment (States II-III, Figure 3b). Physiological environment comprised incubation in complete DMEM at 37°C in a humidified 5% CO_2_ atmosphere to mimic cell culture conditions. In State I (properties), hydrogels were characterized 24 h post-gelation, after reaching an equilibrium of network as confirmed by constant G′ values over time, and subsequently evaluated for network dynamics, viscoelastic properties, and injectability (Sections 3.4-9). State II (physiological environment) was included to assess time-dependent stability and mechanical response of representative formulations (AH-N and DH-N) under static conditions and dynamic loading with medium perfusion in a bioreactor (Sections 3.10). Finally, in State III (cell culture), the biological performance of selected hydrogel formulations was assessed using MSCs and chondrocytes, focusing on cytocompatibility, cell distribution, post-injection cell viability, and chondrocyte shape in three-dimensional environments (Sections 3.11-12).

### 3.4. Viscoelastic properties

The viscoelastic response of hydrogels was evaluated by oscillatory rheology, using amplitude and frequency sweeps to investigate their deformation- and time-dependent mechanical behavior. The amplitude sweep response (Figure 4a) shows G′ exceeding G″ at low strains (≤10%), confirming the formation of solid-like gel networks in all formulations. AH-N, DH-N, and DL-N exhibited an LVR characterized by a strain-independent G′ plateau extending to approximately 20% strain. However, AH-A showed a gradual increase in G′ at low-to-intermediate strains, indicating strain stiffening before the yield point at 50% strain. Beyond the LVR, all hydrogels undergo a transition to liquid-like behavior (G″ > G′), with AH-A showing a higher flow point (∼350% strain) compared to ∼200% strain for the other formulations, highlighting its enhanced resistance to large deformation.

**Figure 4.**
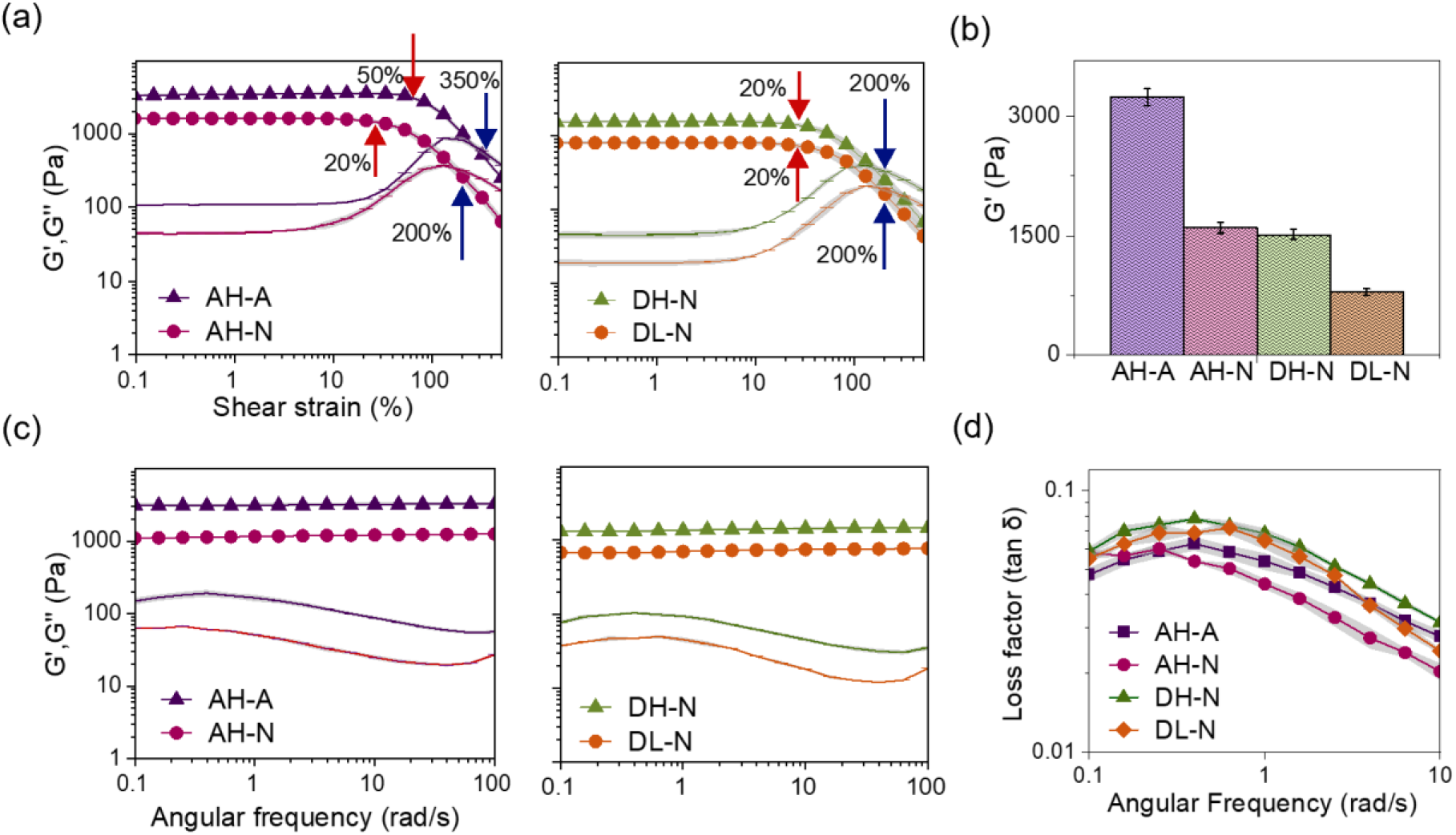
Viscoelastic characterization of hydrogels AH-A, AH-N, DH-N, and DL-N. (a) amplitude sweep measurements showing the LVR, yield (red arrows), and flow (blue arrows) points at frequency of 10 rad·s^-1^, and (b) G′ value within LVR, (c) frequency sweep test at strain 0.2%, (d) loss factor (tan δ) evaluated within the LVR over the frequency range 0.1-10 rad·s^-1^ at a strain of 0.2%. G′ is shown by symbols connected by lines, and G″ is shown by lines only. Rheological measurements were performed in triplicate, with variability shown as shaded regions around the mean curves. Where shading is absent, the standard deviation was negligible.

Consistent with these findings, the G′ values at low strain (0.2%) (Figure 4b), which represent hydrogel stiffness, differ markedly among the formulations. AH-A shows the highest G′ values (3200 ± 100 Pa). AH-N and DH-N exhibit intermediate G′ values (1600 ± 60 Pa and 1500 ± 60 Pa, respectively), despite differences in OxPs backbone, while DL-N presents the lowest stiffness (800 ± 40 Pa). Overall, hydrogel stiffness, follows the order AH-A > AH-N ≈ DH-N > DL-N. These values fall within the typical range reported for acylhydrazone polysaccharide hydrogels (10^2^–10^3^ Pa), where modulus scales with the crosslink density and network architecture.^[5d, 8a]^

Figure 4a reveals that the enhanced stiffness observed in OxA-based hydrogels formed under acidic conditions (AH-A), relative to those prepared under neutral conditions (AH-N), highlights the influence of pH on acylhydrazone network formation and stabilization. In particular, pH-dependent crosslinking kinetics and thermodynamic equilibrium can significantly alter the effective crosslink density of the hydrogel network. The accelerated forward reaction rate promotes a greater fraction of active crosslinks, ultimately resulting in increased network stiffness.^[36]^ In acylhydrazone networks, mildly acidic conditions (∼pH 4.5) accelerate bond formation, as discussed in Section 3.2, while also influencing the thermodynamic equilibrium. These observations are consistent with previous studies demonstrating a direct relationship between bond affinity and the macroscopic mechanical properties of polymer networks.^[3a, 3b, 8b]^ For example, Morgan et al. showed that variations in bond affinity among imine-type dynamic covalent chemistries lead to significant differences in shear modulus, even under identical stoichiometric conditions.^[3a]^ Similarly, Ollier et al. reported pH-dependent mechanical reinforcement in PEG-based dynamic covalent hydrogels crosslinked through reversible boronate ester bonds, where a shift in the binding equilibrium toward the bound state increased the effective crosslink density, resulting in a higher G′.^[36]^ On the other hand, although AH-A and DH-N hydrogels are expected to exhibit comparable effective crosslink densities due to their identical [NH_2_] : [C=O] ratio, AH-A demonstrated a higher G′. This is particularly notable given that acylhydrazone bond formation in OxA-based hydrogels is accelerated under acidic conditions, whereas OxD-based hydrogels exhibit pH-independent gelation behavior. Since both systems were expected to reach similar final network densities under the experimental conditions, the observed difference in mechanical properties is likely not governed by crosslink density alone. Instead, it may originate from structural differences between the OxPs used in this work, consistent with previous reports showing that structural variations among OxPs affect hydrogel viscoelasticity.^[37]^

Figure 4b shows that hydrogel prepared from OxD with lower DO exhibited reduced stiffness compared to hydrogel with higher DO, despite maintaining an identical [NH_2_]: [C=O] of 1 : 2. To preserve this stoichiometric balance, the concentration of Alg-ADH (the component increasing the stiffness) was decreased from 1.3 to 0.6 wt%, while the OxD (the component lowering the stiffness) concentration was increased from 0.6 to 0.8 wt% (Table 1). Consequently, the lower G′ observed for DL-N relative to DH-N (Figure 4b) may be partially attributed to the reduced concentration of Alg-ADH within the network. In addition, Morgan et al. demonstrated that variations in network architecture, particularly in crosslinker valency, defined as the number of reactive functional groups per crosslinking unit, can substantially influence the G′ independently of the thermodynamics of dynamic covalent bond formation.^[3a]^ These findings indicate that hydrogel stiffness is governed not only by bond chemistry, but also by network architecture parameters, i.e., the difference in valency resulting from low DO or the relative ratio of Alg-ADH to OxD within the hydrogel.

Figure 4c presents the frequency-dependent viscoelastic response of the hydrogels. Across the investigated angular frequency range (0.1–100 rad·s^-1^), G′ remains nearly constant while G″ shows a slight increase, consistent with viscoelastic solid-like behavior governed by a stable elastic network. No crossover between G′ and G″ is observed within the accessible frequency window at 37 °C, indicating that the gel-to-sol transition is not reached under these conditions. This behavior is typical for hydrazone- and acylhydrazone-crosslinked hydrogels, where dynamic covalent bond exchange occurs on longer timescales, where lower frequencies are required to capture the gel-to-sol transition.^[5a, 8b]^

To further compare viscoelastic behavior independent of absolute modulus values, the loss factor (tan δ = G″/G′) was evaluated within the LVR, as shown in Figure 4d. All formulations exhibit low tan δ values ranging from approximately 0.02 to 0.08 across the frequency range, confirming the solid-like networks. These values are consistent with those reported for OxA hydrogels crosslinked via ADH ^[6, 14b]^, Alg-ADH hydrogels crosslinked via OxA,^[10b]^ and hyaluronic acid-ADH hydrogels crosslinked via OxHA,^[4a, 10a, 38]^ which typically exhibit tan δ < 0.1 under small-amplitude oscillatory shear.

### 3.5. Network recovery

The self-recovery behavior of the acylhydrazone hydrogels was evaluated using two deformation protocols, i.e., progressive strain–amplitude^[25]^ and cyclic step-strain.^[26]^ Although both protocols apply a maximum strain of 500% followed by recovery at 0.2% strain, they differ in their deformation history. In the progressive strain–amplitude test, the strain is gradually increased from 0.1% to 500% over approximately 5 min, resulting in continuous network deformation. In contrast, the cyclic step-strain test applies an abrupt transition from 0.2% to 500% strain within approximately 1 s, imposing an instantaneous large deformation.

Figure 5a presents a progressive strain–amplitude recovery experiment providing the normalized viscoelastic moduli (G’ and G’’) as a function of time. These data illustrate the viscoelastic response during progressive deformation and the subsequent recovery of the hydrogel network, as reflected by the recovery of the G′ after the strain is returned to 0.2%. The initial stage of this experiment follows the same gradual strain-ramp protocol as the amplitude sweep shown in Figure 4a, in which the strain is continuously increased from 0.1% to 500%. Following this large deformation, all formulations exhibited a characteristic two-stage recovery after returning to 0.2% strain. An initial rapid recovery restored G′ to approximately 90% of its original value, followed by a slower recovery phase associated with progressive reformation of the dynamic covalent network, ultimately reaching full recovery (∼100%) within 15–30 min. AH-A recovered faster than AH-N, requiring 15 min and 30 min, respectively, to recover after deformation, whereas DH-N and DL-N exhibited comparable recovery times of 20–23 min, similar to AH-A (Figure 5a). The faster recovery of AH-A relative to AH-N is attributed to the accelerated kinetics of acylhydrazone bond formation under mildly acidic conditions (Figure 2c). In addition, the comparable recovery times of AH-A, DH-N, and DL-N correlate with their similarly rapid gelation kinetics, indicating fast bond formation within these hydrogel networks (Figure 2c and Figure S3).

**Figure 5.**
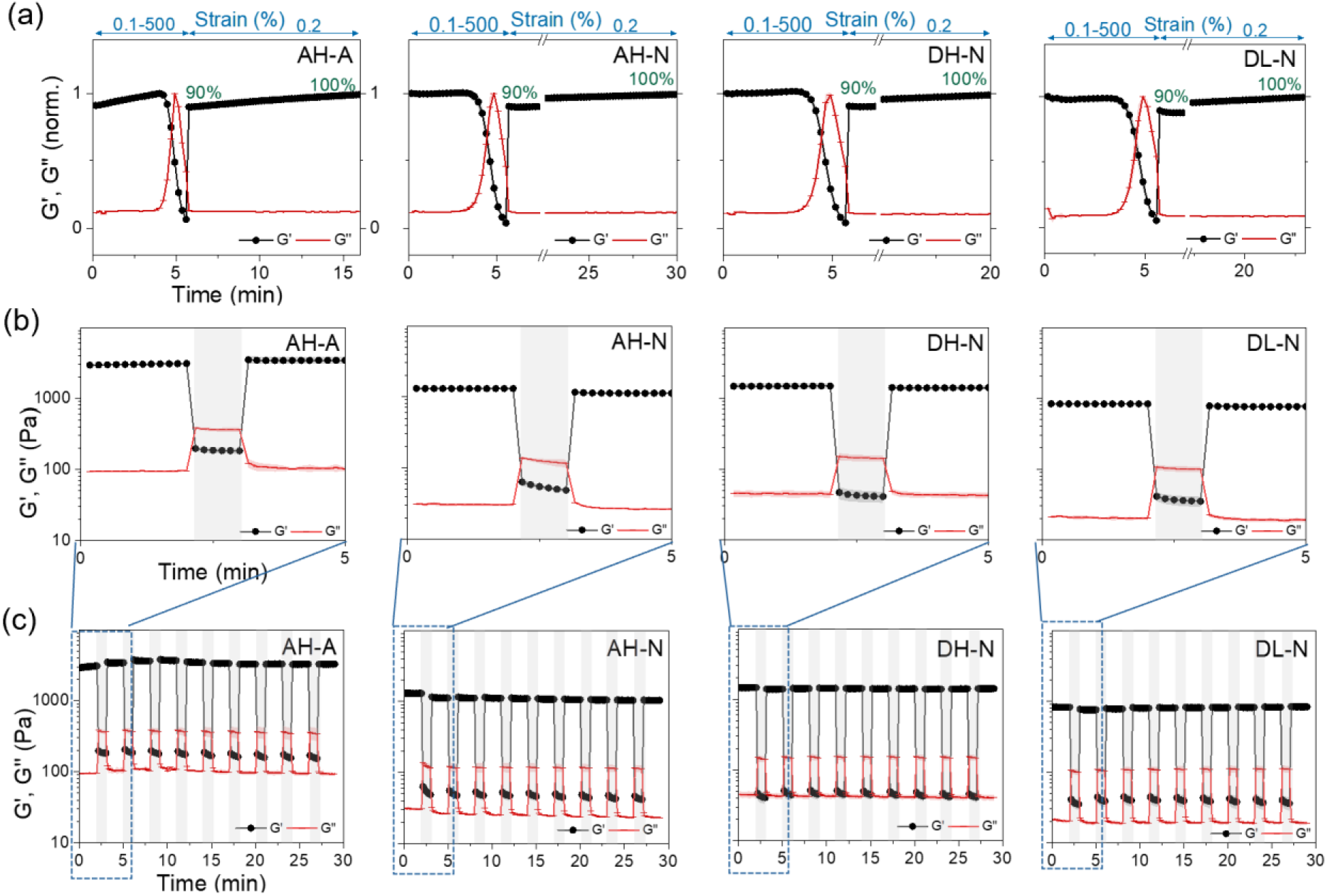
Network recovery of acylhydrazone hydrogels. (a) Progressive strain– amplitude tests. G′ was normalized to its maximum values to demonstrate full recovery at G′ = 1. Cyclic step-strain tests for hydrogels subjected to alternating low (0.2%) and high (500%, highlighted area) strain, (b) magnified view of the first cycle, and (c) over ten cycles. Rheological measurements were performed in triplicate, with variability shown as shaded regions around the mean curves. Where shading is absent, the standard deviation was negligible.

Figures 5b and 5c show cyclic step-strain measurements, where abrupt alternating low- and high-strain intervals were applied. During the first cycle (Figure 5b), AH-A exhibited an over-recovery response, reaching 120 ± 15% of its initial G′, whereas AH-N recovered to 90 ± 5%, and DH-N and DL-N reached 95 ± 10% and 95 ± 5%, respectively. After ten cycles (Figure 5c), AH-A, AH-N, DH-N, and DL-N recovered to 110 ± 4%, 80 ± 10%, 97± 12%, and 100 ± 4% of their initial G′, respectively, indicating effective recovery under repeated deformation. Although the gelation kinetics of DH-N and DL-N were relatively rapid and comparable to AH-A (Figure 2c and Figure S3), the over-recovery of AH-A apparently reflects additional network rearrangement during the post-deformation recovery process that is absent in recovery of OxD-based hydrogels.

### 3.6. Compressive mechanical properties

The compressive mechanical properties of the acylhydrazone hydrogels were evaluated by unconfined uniaxial compression. Figure 6a presents representative compressive stress–strain curves for AH-A, AH-N, DH-N, and DL-N hydrogels, with the corresponding compressive moduli shown in Figure 6b. All formulations exhibit a nonlinear stress–strain response consisting of an initial low-strain regime (0–10%) followed by progressive strain stiffening between approximately 30–40% strain and failure at strains of approximately 50–60% at a failure stress in the range of 40–65 kPa. This behavior is characteristic of soft viscoelastic polymer networks, in which network densification, polymer chain alignment, and restricted segmental mobility contribute to increased stiffness under compression.^[39]^ Figure 6b shows that AH-A exhibited the highest compressive modulus, reaching 20 ± 3 kPa. AH-N and DH-N exhibited comparable compressive moduli of 11 ± 3 kPa and 12 ± 2 kPa, respectively, whereas DL-N showed the lowest compressive modulus at 9 ± 1 kPa. Overall, the hydrogels follow the order of compressive modulus AH-A > DH-N ≈ AH-N > DL-N, while exhibiting comparable failure strains (50–60%). This suggests that the formulations primarily govern hydrogel stiffness rather than the hydrogel failure strain.

**Figure 6.**
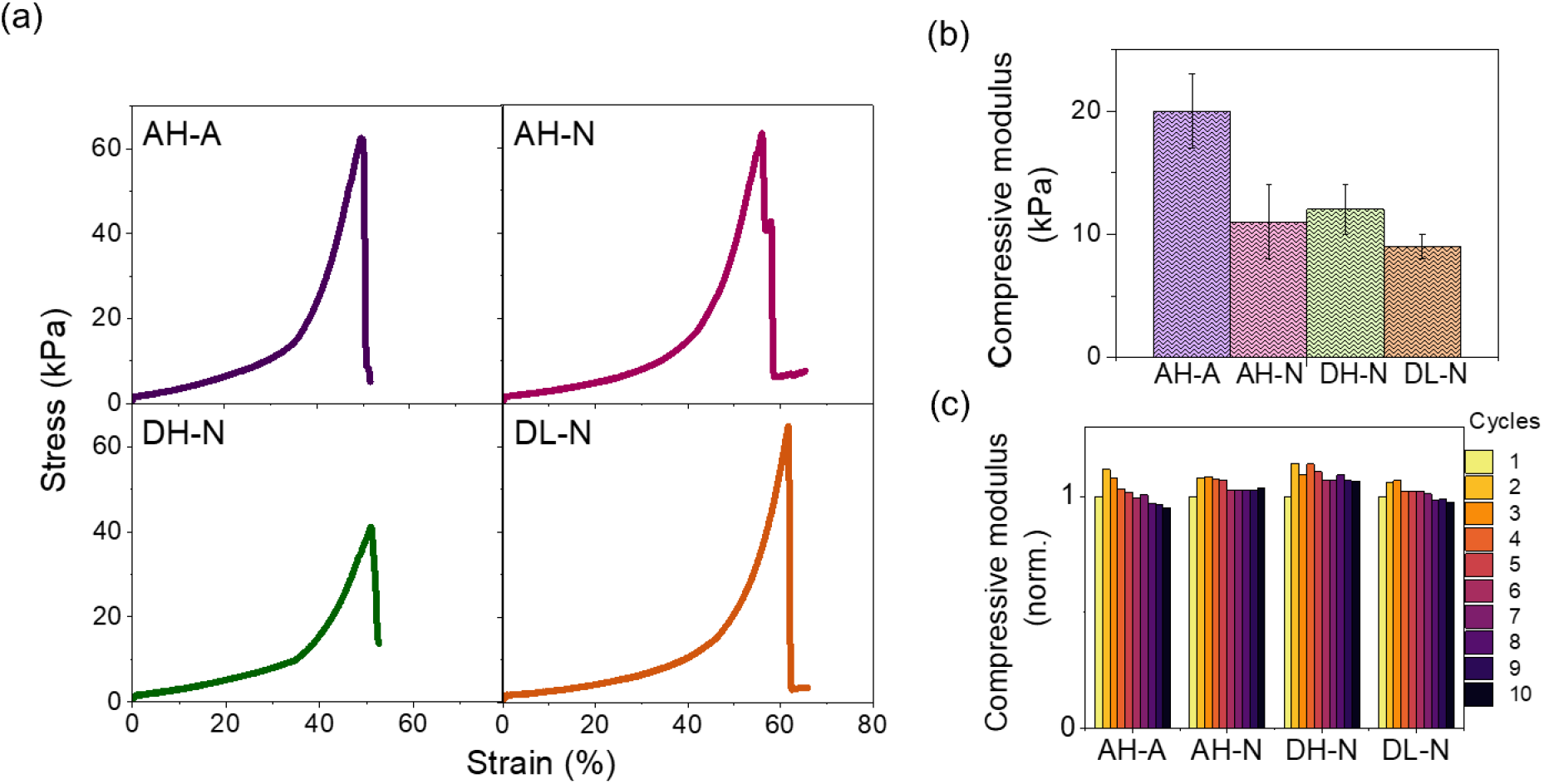
Compressive mechanical properties of AH-A, AH-N, DH-N, and DL-N acylhydrazone hydrogels. (a) Representative unconfined compressive stress–strain curves recorded up to hydrogel failure, defined by the maximum of the curve. (b) Compressive moduli determined from stress-strain curves. (c) Cyclic unconfined compression to a strain of 20% resulting in compressive modulus normalized to the initial compressive modulus.

Figure 6c shows the cyclic compressive behavior of the hydrogels, where the normalized compressive modulus is plotted over ten loading–unloading cycles at a strain of 20%. For all formulations, the first loading cycle exhibits a lower modulus compared to the second cycle. This phenomenon is reported in soft materials and biological tissues, where repeated loading leads to a transition toward a steady-state mechanical response through a combination of viscoelastic relaxation, fluid redistribution, and microstructural rearrangement within the network.^[40]^ After the second cycle, AH-A shows a slight reduction in modulus with increasing cycle number, suggesting minor cyclic softening, while AH-N, DH-N, and DL-N maintain nearly constant responses. The absence of significant modulus decay indicates that the hydrogels possess robust structural integrity and fatigue resistance under cyclic loading. At a glance this stiffening seen for all the hydrogels demonstrates different behavior compared to the strain-step, where the stiffening was only seen for AH-A sample. The reason for the difference between these two experiments is seen in different deformation modes and loading protocols.

### 3.7. Viscoelastic stress relaxation

Stress relaxation in dynamic acylhydrazone hydrogels arises from the coexistence of multiple relaxation processes, including polymer-chain rearrangement, reversible crosslink dissociation/reformation, and network-level structural reorganization, which together determine the time-dependent viscoelastic response of the material,^[4b, 5d, 41]^ as schematically illustrated in Figure 7a. To investigate these relaxation kinetics or behavior under different mechanical environments, stress relaxation experiments were performed under compression and shear deformation at a constant strain of 15%, selected within the initial low-strain compressive region and within the LVR of all hydrogel formulations. Compressive deformation primarily reflects the bulk structural stability, load-bearing capability, and network rearrangement, whereas shear deformation is more sensitive to chain mobility, transient network rearrangements, and dynamic crosslink-mediated relaxation processes.^[8b, 42]^ As shown in Figures 7b and 7c, all hydrogels exhibited stress relaxation behavior, with the normalized stress profiles revealing formulation-dependent differences in relaxation kinetics under both deformation modes.

**Figure 7.**
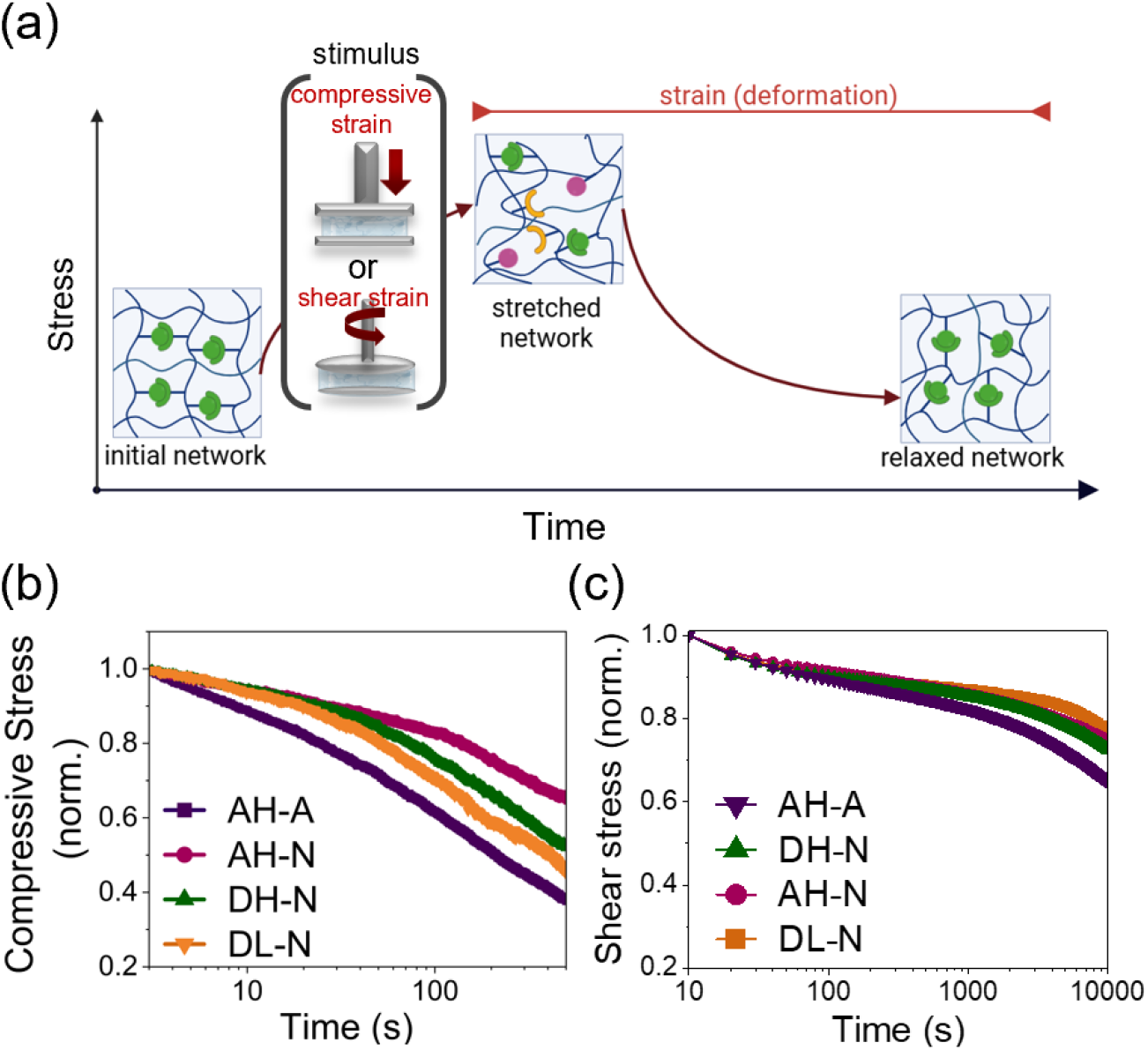
Stress-relaxation behavior of AH-A, AH-N, DH-N, and DL-N acylhydrazone hydrogels. (a) Schematic illustration of network rearrangement enabled by reversible crosslinks under applied shear or compressive deformation. (b) Compressive stress at 15% strain and (c) shear stress relaxation profile at 15% strain.

Among the investigated hydrogels, AH-A displayed the fastest stress relaxation in both compression (Figure 7b) and shear starin (Figure 7c), indicating enhanced molecular mobility and more efficient stress dissipation, consistent with higher exchange dynamic of acylhydrazone bonds. In contrast, AH-N exhibited a substantially slower relaxation profile compared to AH-A, reflecting the presence of more slowly forming crosslinks and reduced network dynamics. This observation is in agreement with previous reports in dynamic supramolecular hydrogels demonstrating that networks formed through rapidly associating crosslinks generally exhibit faster stress relaxation than those formed through slower dynamic interactions.^[43]^

Although AH-A and DH-N, and DL-N exhibited comparable rapid gelation behavior associated with fast forward acylhydrazone formation, AH-A demonstrated faster stress relaxation under both deformation modes. This finding indicates that the rate of network formation does not directly determine the relaxation behavior of the resulting hydrogel network with different OxPs. Instead, the relaxation response is governed by the dynamic characteristics of the formed network, including rate of bond dissociation, crosslink accessbility, and polymer-chain mobility. Therefore, differences between AH-A and DH-N and DL-N likely arise from variations in the molecular arrangement and accessibility of dynamic crosslinking sites after network formation.

Similarly, although AH-N exhibited slower gelation kinetics due to a lower forward acylhydrazone formation rate, its stress relaxation behavior under shear deformation became comparable to that of DH-N after network formation. This observation further demonstrates that the influence of the crosslinking kinetics decreases after the dynamic covalent network is formed, and the subsequent time-dependent mechanical properties are mainly controlled by the reversibility of acylhydrazone bonds and the structural characteristics of the polymer network. Under compression, however, AH-N exhibited slower relaxation than DH-N, indicating that bulk network organization and macroscopic load-transfer contribute additionally to stress dissipation under compressive deformation. Overall, these results demonstrate that the forward acylhydrazone formation rate primarily regulates gelation kinetics, whereas the relaxation behavior is governed by the interplay between exchange dynamic (particularly the rate of dissociation) and polymer network architecture. The chemical structure and oxidation characteristics of the OxPs therefore provide a means to independently tune network formation and time-dependent mechanical properties.

The shear stress relaxation data were further analyzed using a two-mode Maxwell– Weichert model. All formulations revealed a characteristic biphasic decay for all formulations. An initial rapid stress relaxation occurred within the first ∼100–300 s, followed by a slower decay over longer timescales. This response is expected for dynamic covalent hydrogels and reflects the coexistence of two dominant relaxation mechanisms: fast polymer network rearrangement and lower rate of hydrazone exchange.^[4b, 5d]^ Such dual-mode relaxation behavior is consistent with the viscoelastic response of many soft biological tissues,^[44]^ where rapid energy dissipation is coupled with long-term structural remodeling. The relaxation profiles in Figure 7c were well-described by a bi-exponential Maxwell–Weichert model consisting of two Maxwell elements in parallel.^[5d]^ The agreement between the experimental data and the model fit (R^2^ > 0.99, Table S7, Figure S5, Supplementary Information) confirms that stress relaxation in these systems is governed by the two dominant relaxation modes. Across all hydrogel formulations, the fast relaxation timescale τ_1_ ranged from 37 to 180 s, whereas the slow relaxation timescale τ_2_ ranged from 6,000 to 12,000 s. The relative stress contribution of the fast relaxation mode, σ1, ranged from 0.4 to 0.6, while that of the slow relaxation mode, σ_2_, ranged from 0.4 to 0.6 (Table S7).

To verify whether the observed stress relaxation behavior was representative across different deformation amplitudes, shear stress relaxation was subsequently evaluated over a strain range of 5–100%, spanning both the LVR and non-LVR while remaining below the flow point. This analysis was performed to determine whether the relaxation mechanism is governed primarily by dynamic exchange of acylhydrazone bonds or is influenced by strain-induced alterations in the network structure. Figure 8a demonstrates representative shear stress–time curves corresponding to shear stress as a function of time, and Figure 8b shows the maximum stress as a function of strain for all formulations. In all hydrogels, the initial stress increased with increasing strain amplitude. The absence of an abrupt decrease in stress (Figure 8b) indicates that network failure did not occur within the investigated strain range from 5 to 100%. Differences in initial stress among formulations (Figures 8a and 8b) correspond to their stiffness, shown as G’ in Figure 4b and compressive modulus in Figure 6b. AH-A shows the steepest increase that is consistent with its highest G’ and compressive modulus (Figure 8b). In this regard, the relative order of formulations was maintained across the entire strain range, with AH-A > AH-N ≈ DH-N >DL-N.

**Figure 8.**
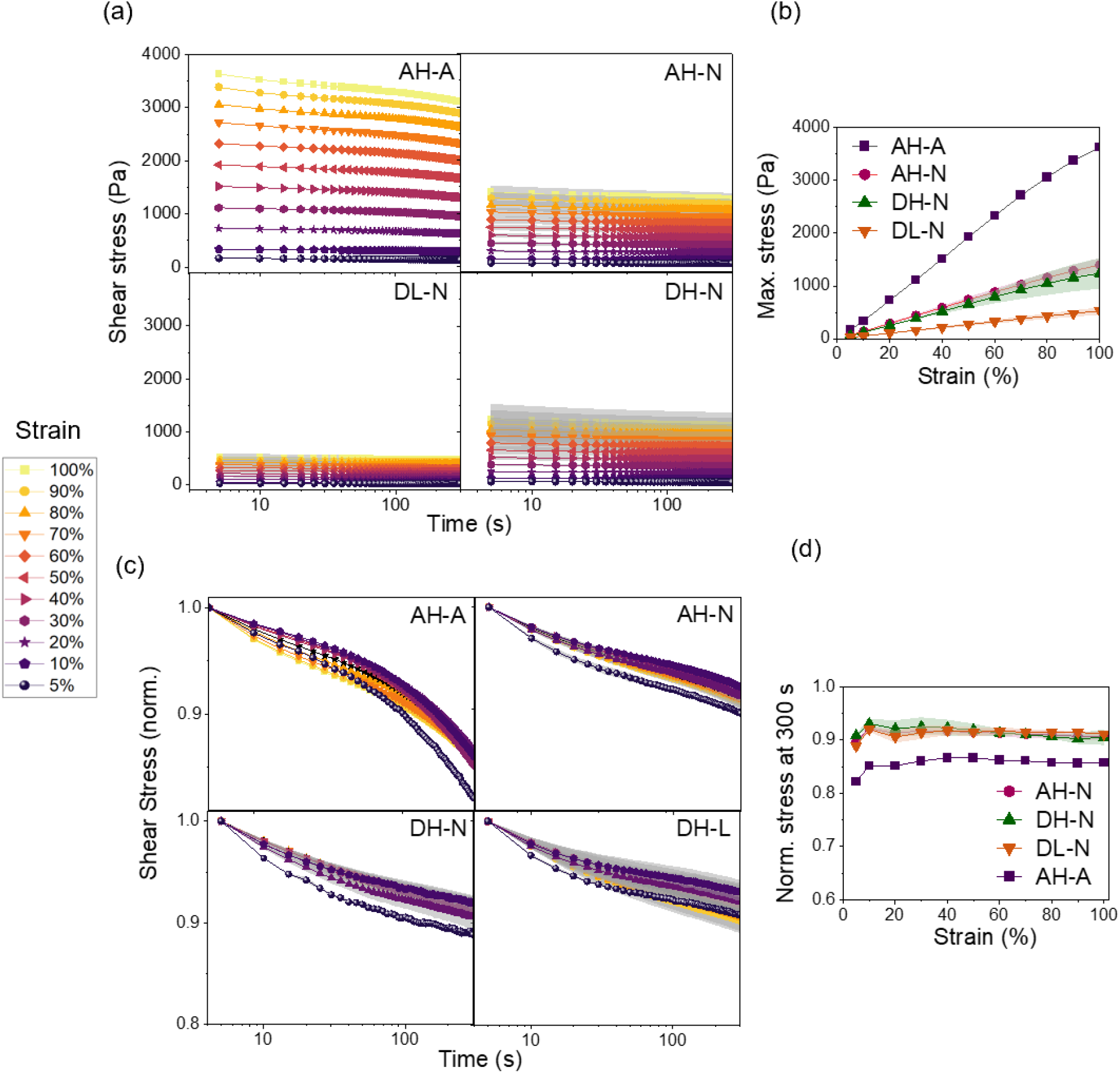
Stress-relaxation behavior of AH-A, AH-N, DH-N, and DL-N acylhydrazone hydrogels. (a) Shear stress–time profiles obtained from step-strain relaxation experiments at strain amplitudes ranging from 5% to 100%. (b) Maximum shear stress as a function of applied strain. (c) Corresponding shear stress normalized to the initial shear stress as a function of time. (d) Normalized shear stress remaining after 300 s as a function of strain.

Figure 8c shows the corresponding normalized shear stress that, upon application of a constant strain, all hydrogels exhibited a gradual decay over the 300 s relaxation period within both LVR and beyond. Interestingly, although the absolute stress magnitudes vary significantly (Figure 8a) in each formulation, the corresponding normalized stress relaxation curves (Figure 8c) reveal similar relaxation kinetics. Figure 8d shows that no pronounced strain dependence was observed across the investigated strain range. Comparable stress relaxation observed in LVR and beyond indicates that dynamic exchange of acylhydrazone bonds and network rearrangement occur under deformation conditions in which the network structure remains mechanically intact.

### 3.8. Injectability of acylhydrazone hydrogels

For dynamic hydrogels intended for minimally invasive surgical administration, injectability reflects the ability of the dynamic network to undergo reversible disruption during extrusion, enabling flow through a syringe and recovery after removal of the applied stress. The injectability of the acylhydrazone hydrogels was evaluated using combined syringe extrusion experiments to assess injection force and shear-thinning/self-healing behavior (Figure 9a–d, Figure S6, and Video S1-4, Supplementary Information) and to correlate dynamic network structure with flow and recovery behavior.

**Figure 9.**
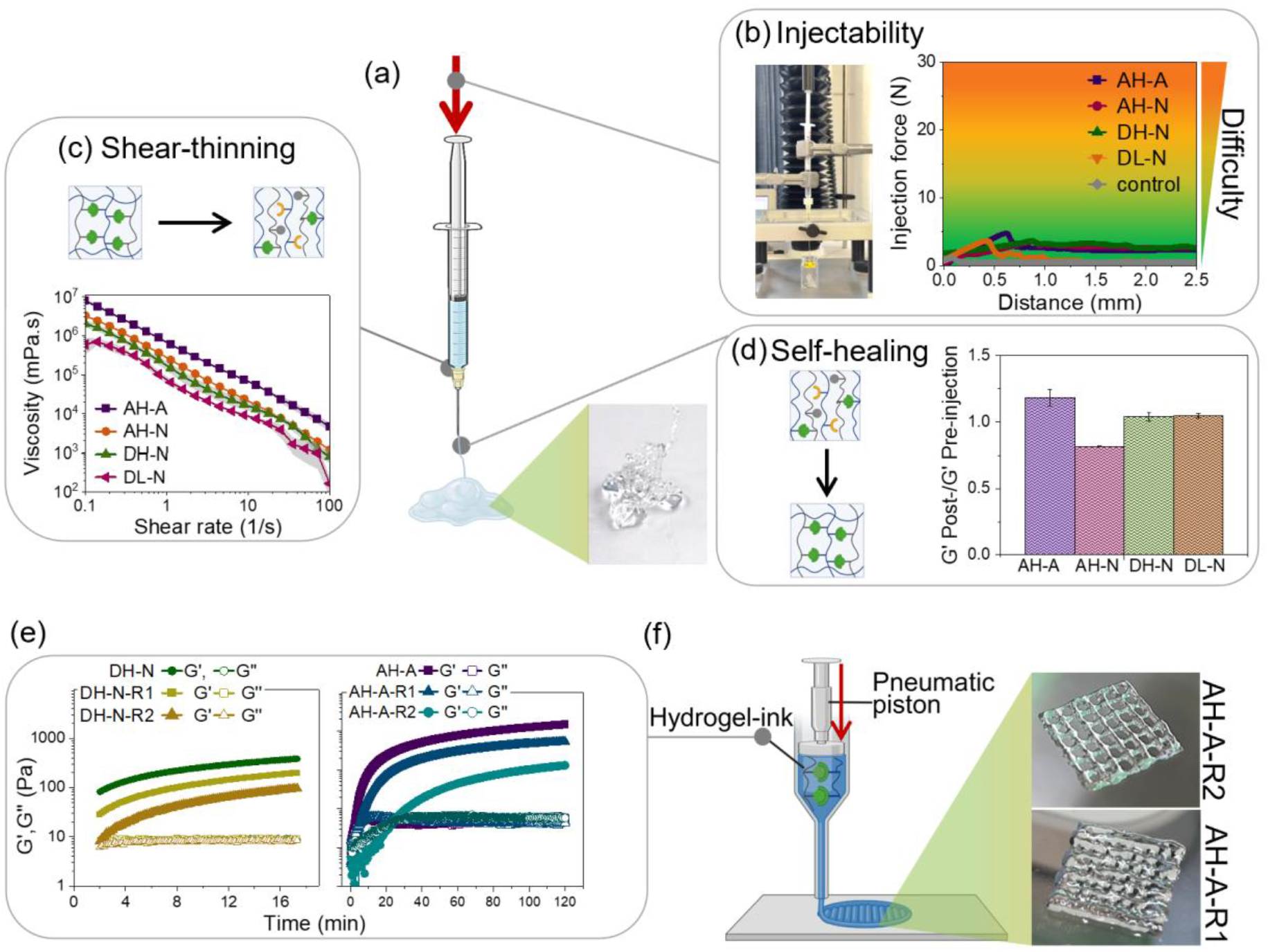
Injectability and printability of acylhydrazone hydrogels. (a) Schematic illustration of syringe injection process, together with a representative image of the extruded hydrogel. (b) Injection force curves recorded during syringe extrusion. (c) Viscosity as a function of shear rate. (d) Recovery of G’ modulus after extrusion, expressed as the ratio of G’ determined post- and pre-injection, respectively. (e) Time-sweep rheological measurements for hydrogels. (f) Schematic illustration of extrusion-based bioprinting together with representative images of printed hydrogel constructs.

Figure S6 and Video S1-4 demonstrate that all hydrogels were readily extruded through a 21 G needle without clogging, fracture, or phase separation while maintaining their structural integrity after extrusion, as shown in Figure 9a. To evaluate the applied force required for surgical administration of injectable hydrogels, the injection force profiles were recorded as a function of plunger displacement during the syringe injection of hydrogels through a 21 G needle. Figure 9b shows that the measured extrusion force for all hydrogels remained substantially below the accepted manual injection threshold of 30 N reported for surgical applications.^[30]^ The measured extrusion forces are consistent with previously reported data by Leo et al. for a 1.5 wt% acylhydrazone-crosslinked hyaluronic acid hydrogel extruded through 18–25 G needles.^[10a]^

Figure 9c shows that in the steady-shear rheological profiles, all formulations exhibited pronounced shear-thinning behavior, with the apparent viscosity decreasing by several orders of magnitude as the shear rate increased from 0.1 to 100 s^-1^. Such shear-thinning behavior is a key requirement for injectable biomaterials, as it enables facile extrusion through high-gauge needles while maintaining structural integrity under low shear conditions. This behavior is consistent with previously reported acylhydrazone-based polysaccharide hydrogels, which combine shear-induced network disruption with rapid bond reformation and self-healing after cessation of flow.^[6a, 9a, 11, 15, 45]^ Among the tested formulations, AH-A exhibited the highest and DL-N the lowest zero-shear viscosity, which is in agreement with the trends observed in compressive modulus and G′ of the hydrogels, as shown in Figures 4b and 6b.

Following extrusion, injectable hydrogels need to rapidly recover their network through self-healing to maintain mechanical integrity at the delivery site. This behavior was quantified by the ratio of post-injection to pre-injection G’ (G’_post_/G’_pre_). Figure 9d schematically depicts the recovery mechanism of a disrupted polymer network through the reformation of covalent crosslinks. This behavior aligns with previously reported syringe injectability of OxA- and OxD-based acylhydrazone hydrogels crosslinked using ADH, which exhibit shear-thinning during extrusion followed by rapid network recovery and self-healing.^[6a, 10b, 15]^ While earlier studies demonstrated injectability of individual formulations, the present work establishes how OxPs chemistry and gelation kinetics influence injectability through their effects on the rheological response of the hydrogel network.

Quantitatively, AH-A demonstrated the highest recovery of approximately 1.2-fold, indicating the full restoration and even slight reinforcement of the network, likely due to post-shear structural rearrangement. In contrast, AH-N exhibited the most reduced recovery of around 0.8-fold, suggesting incomplete network recovery at the measured time point (approximately 30 min after extrusion), potentially due to lower dyanmic exchange. The DH-N and DL-N samples display intermediate performance, with G′ ratios of 1.0–1.05, indicating the complete recovery of viscoelastic properties. This recovery behavior of hydrogels is consistent with their cyclic step-strain experiments (Figures 5b and 5c), where AH-A exhibited over-recovery, DH-N and DL-N recovered completely, and AH-N showed slightly reduced recovery. Additionally, frequency-sweep measurements on post-injected hydrogels (Figure S7, Supplementary Information) revealed no significant differences in viscoelastic profiles compared with pre-injection samples (Figure 4b), confirming that shear loading does not induce permanent structural damage. The persistence of a G’-dominated response further confirms the rapid reformation of the elastic network upon removal of shear stress.

### 3.9. Extrusion printability of acylhydrazone hydrogel inks

Hydrogels with fast gelation kinetics (Figure 2c) and the most effective recovery (Figure 5 and Figure 9d), AH-A and DH-N, were selected for extrusion printability. The ability of these hydrogels to undergo continuous flow and to form stable, self-supporting filaments upon deposition was evaluated using an extrusion-based bioprinting setup. Although both hydrogels exhibited successful injectability, they failed to generate continuous printed strands under the printing conditions as defined in the experimental section (Section 2.8). Instead, they produced discontinuous filaments and irregular deposition patterns, indicating an insufficient stability of the extruded filament. This discrepancy highlights that injectability alone is not a reliable predictor of printability, as extrusion bioprinting requires a finely balanced interplay between shear-induced flow and rapid structural recovery to preserve filament integrity after deposition.^[46]^

Here we focused on the role of crosslinking stoichiometry on printability as a design parameter to access a printable processing window in relation to the gelation kinetics. Accordingly, the [NH2] : [C=O] molar ratio was changed from 1 : 2, corresponding to the main hydrogel library (Table 1), to 1 : 1 (R1) and 2 : 1 (R2) in AH-A-R and DH-N-R formulations (Table 2) to modulate gelation kinetics. Figure 9e showed that increasing the [NH2] to [C=O] ratio reduced G′, while G″ remained unchanged. These G′ values indicate a reduction in effective crosslink density and a delayed evolution of the elastic network.

Based on these rheological insights, hydrogel inks were printed at a defined stage of network development, characterized by the evolution of G′ over time. The extrusion was performed for AH-A-R2 and AH-A-R1, near their crossover point approximately 20 min and 5 min post-mixing (Figure 9e), respectively. At this point, the AH-A-R2 exhibited an optimal balance between flowability and structural integrity, enabling continuous filament formation during deposition. As shown in Figure 9f, AH-A-R2 produced uniform, continuous filaments, and the printed constructs maintained high shape fidelity after deposition, confirming successful extrusion-based printability. However, AH-A-R1 exhibited partial filament collapse and deformation after deposition, resulting in lower structural fidelity and less defined pore architecture. This behavior is likely associated with faster network development (faster increase of G’) of the AH-A-R1 formulation, which limited smooth filament spreading and stabilization during layer-by-layer deposition. OxD-based formulations did not form continuous filaments, attributable to their faster gelation kinetics and reduced processing window, which limit stable extrusion under identical printing conditions.

### 3.10. Time-dependent stability and mechanical response under physiological environment

The stability of the hydrogels and their dependence on the DO of the OxPs, crosslinking conditions, and incubation mode were evaluated. Hydrogel stability was assessed in complete DMEM at 37 °C under 5% CO2, as acylhydrazone bonds are known to exhibit different stability in biological media compared with buffered solutions.^[34]^ Figure 10a presents representative images of MSCs-laden hydrogels cultured under static conditions for up to 90 days. During incubation, the hydrogels exhibited a gradual increase in volume, followed by progressive loss of macroscopic integrity. Complete disappearance of the visible macroscopic structure of hydrogels was defined as the “GONE” state. Overall, hydrogels formed via high-DO OxPs (DH-N, AH-N, and AH-A) displayed substantially longer persistence than those prepared from low-DO OxPs (DL-N, AL-N, and AL-A). Among the low-DO formulations, DL-N maintained its macroscopic structure for 40 days, whereas AL-A and AL-N reached the GONE state after 21 and 14 days, respectively. In contrast, DH-N retained its macroscopic structure throughout the entire 90-day observation period, while both AH-N and AH-A persisted for 60 days.

**Figure 10.**
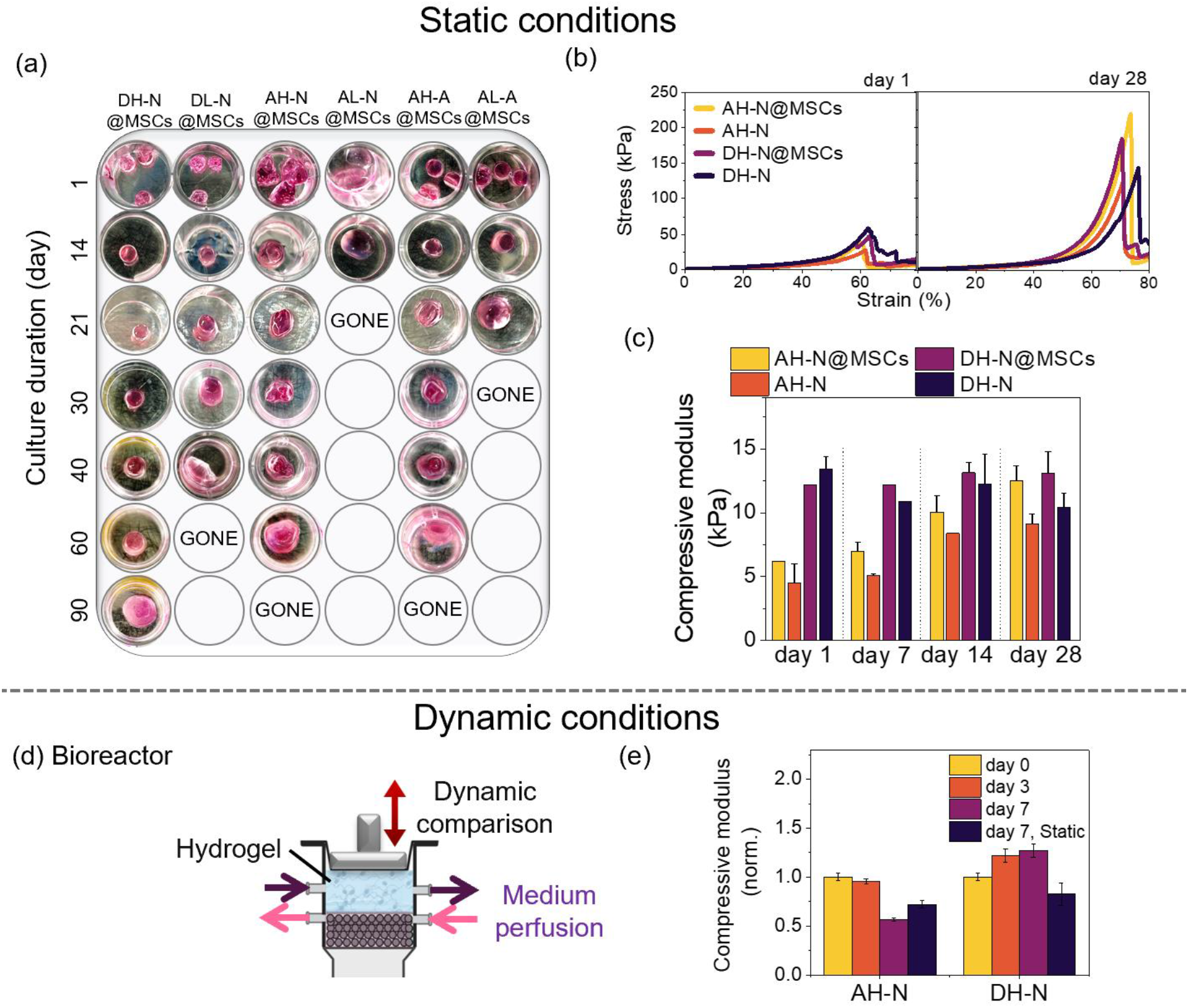
Comparative analysis of static and dynamic conditions for hydrogels with and without laden MSCs. (Top) Static conditions in a standard multi-well plate. (a) Representative images of the hydrogels show variability in shape and sample loss (marked "GONE"). (b) Stress–strain curves for hydrogels and (c) corresponding compressive modulus of cell-laden and cell-free AH-N and DH-N hydrogels during static culture. (Bottom) Dynamic incubation system utilizing a bioreactor setup with dynamic compression and medium perfusion. (d) The schematic depicts a bioreactor with a medium-perfusion system and mechanical stimulation. (e) Compressive modulus of hydrogels exposed to the dynamic conditions in bioreactor compared to static controls. The dynamic compressive modulus was normalized to the corresponding day 0 value (AH-N: 7 ± 0.3 kPa; DH-N: 4 ± 0.1 kPa).

Interestingly, no noticeable difference in stability was observed between AH-N and AH-A, indicating that once the network is formed and exposed to the physiological environment, the initial crosslinking conditions have only a limited influence on stability. The acidic environment of AH-A is expected to be rapidly neutralized upon immersion in DMEM, resulting in pH conditions comparable to those of AH-N. Consequently, both hydrogels are expected to exhibit similar behavior after incubation in complete DMEM. In contrast, among the low-DO formulations, faster gelation contributed to improved hydrogel stability. Representative images obtained on day 1 show that AL-N formed a poorly defined construct, whereas DL-N and AL-A rapidly generated well-defined disc-shaped hydrogels, which subsequently exhibited longer persistence during culture. The prolonged persistence of the high-DO hydrogels is likely attributable to the higher availability of aldehyde groups compared to the low-DO formulations, resulting in improved network connectivity. A previous study investigated the stability of acylhydrazone hydrogels prepared from hydrazide-functionalized gelatin crosslinked with oxidized hyaluronic acid (OxHA) or OxA during 21 days of incubation in DMEM. The OxHA-based hydrogels progressively swelled and lost their macroscopic shape, whereas the OxA-based hydrogels maintained their structural integrity throughout the incubation period.^[47]^

To determine whether the stability was accompanied by changes in mechanical properties, compressive behavior was evaluated in both MSC-laden (AH-N@MSCs and DH-N@MSCs) and cell-free (AH-N and DH-N) hydrogels cultured under static conditions for 28 days. AH-N and DH-N were selected because they combined high stability with cytocompatible neutral gelation conditions. Stress–strain profiles in Figure 10b show that the failure stress of hydrogels increased with culture time, rising from approximately 50 kPa on day 1 to nearly 200 kPa on day 28, irrespective of cell presence. Figure 10c further demonstrates distinct mechanical evolution between the two hydrogel systems. AH-N and AH-N@MSCs exhibited a gradual increase in compressive modulus from day 1 to day 28, whereas DH-N and DH-N@MSCs maintained nearly constant moduli throughout the culture period. These observations suggest that the mechanical stability of the hydrogels is governed not only by structural stability but also by the kinetics of dynamic acylhydrazone network formation. The gradual stiffening of AH-N could be consistent with and network rearrangement and maturation after gelation, reflecting the slower acylhydrazone formation kinetics of this system. In contrast, the rapid bond formation in DH-N produces a mechanically mature network shortly after gelation, resulting in stable mechanical properties during prolonged static culture. Importantly, the mechanical performance of the hydrogels was preserved in the presence of encapsulated MSCs. These characteristics support the suitability of the hydrogels for long-term three-dimensional MSCs culture by providing sustained mechanical performance throughout the culture period.

The mechanical behavior of the dynamic covalent network in AH-N and DH-N hydrogels was further investigated under continuous medium perfusion combined with cyclic compressive loading in bioreactor (Figure 10d). Compared with static conditions, this environment imposes repeated mechanical disturbance while continuously exchanging the surrounding medium, thereby providing a more demanding assessment of the ability of reversible acylhydrazone bonds to dissociate and reform under physiologically relevant conditions. In addition, dynamic compression induces forced advection of the culture medium inside the hydrogels.^[29]^ As shown in Figure 10e, AH-N and DH-N exhibited markedly different responses under dynamic conditions. AH-N underwent an approximately 50% reduction in compressive modulus after 7 days, whereas DH-N displayed an increase in modulus of approximately 30% over the same period. Notably, this behavior contrasts with the static control results (Figure 10c), where DH-N remained mechanically unchanged while AH-N gradually stiffened. These findings demonstrate that static and dynamic conditions assess different aspects of network behavior. Static conditions primarily reflect network maturation after gelation, whereas dynamic conditions evaluate the ability of the dynamic acylhydrazone network to continuously reorganize under repeated mechanical loading and media perfusion. The enhanced mechanical performance of DH-N under bioreactor conditions is likely a consequence of its higher dynamic acylhydrazone exchange, which enable efficient dissociation and reformation of crosslinks during cyclic compression in the presence of medium continue perfusion and forced advection. This continuous exchange facilitates reformation of load-bearing connections, allowing the hydrogel to maintain and even enhance its mechanical integrity through network rearrangement despite repeated deformation. In contrast, the lower dynamic acylhydrazone exchange of AH-N limit efficient network reorganization under the same conditions, leading to progressive mechanical weakening (Figure 10e).

### 3.11. Cell-laden hydrogels

To evaluate the performance of the hydrogels for cell culture, MSCs and chondrocytes were encapsulated within the hydrogels. MSCs are commonly used to assess dynamic hydrogels due to their mechanosensitivity to matrix stiffness and stress relaxation behavior.^[2b, 48]^ Chondrocytes were used as a complementary cell model because their shape and function are regulated by cell–matrix interactions mediated by the viscoelastic properties of the hydrogel network.^[5b, 5c, 7]^ Importantly, chondrocytes can also sense and respond to mechanical cues through adhesion-independent mechanotransduction, in which cell volume confinement within three-dimensional environments regulates cellular behavior independently of adhesive ligands.^[49]^ In addition, chondrocytes have been shown to maintain viability and phenotype in non-adhesive hydrogel networks, such as alginate, without the incorporation of cell-adhesive motifs, whereas MSCs behavior is more dependent on adhesive ligands that mediate cell–matrix interactions.^[2b, 6a]^ Together, these cell models enable a comprehensive evaluation of cytocompatibility and the capacity of the dynamic hydrogels to support cellular behavior under physiologically relevant conditions. Accordingly, key biological and material-related parameters were assessed, including cell viability, spatial distribution, cell morphology, and the effect of the injection process on post-injection cell viability.

#### 3.11.1. MSCs-laden hydrogels

Acylhydrazone hydrogels were investigated under two processing conditions (Figure 11a): MSCs-laden hydrogels were transferred into culture plates by a direct casting using an insulin syringe without a needle to evaluate cytocompatibility of the hydrogel precursors containing aldehyde- and hydrazide-functionalized polymers, the in situ acylhydrazone crosslinking process, and the resulting three-dimensional hydrogel network, which should provide a porous and permeable environment for nutrient and oxygen transport. In parallel, MSCs-laden hydrogels were extruded through a 21 G needle to investigate the effect of the injection process on cell viability and distribution.

**Figure 11.**
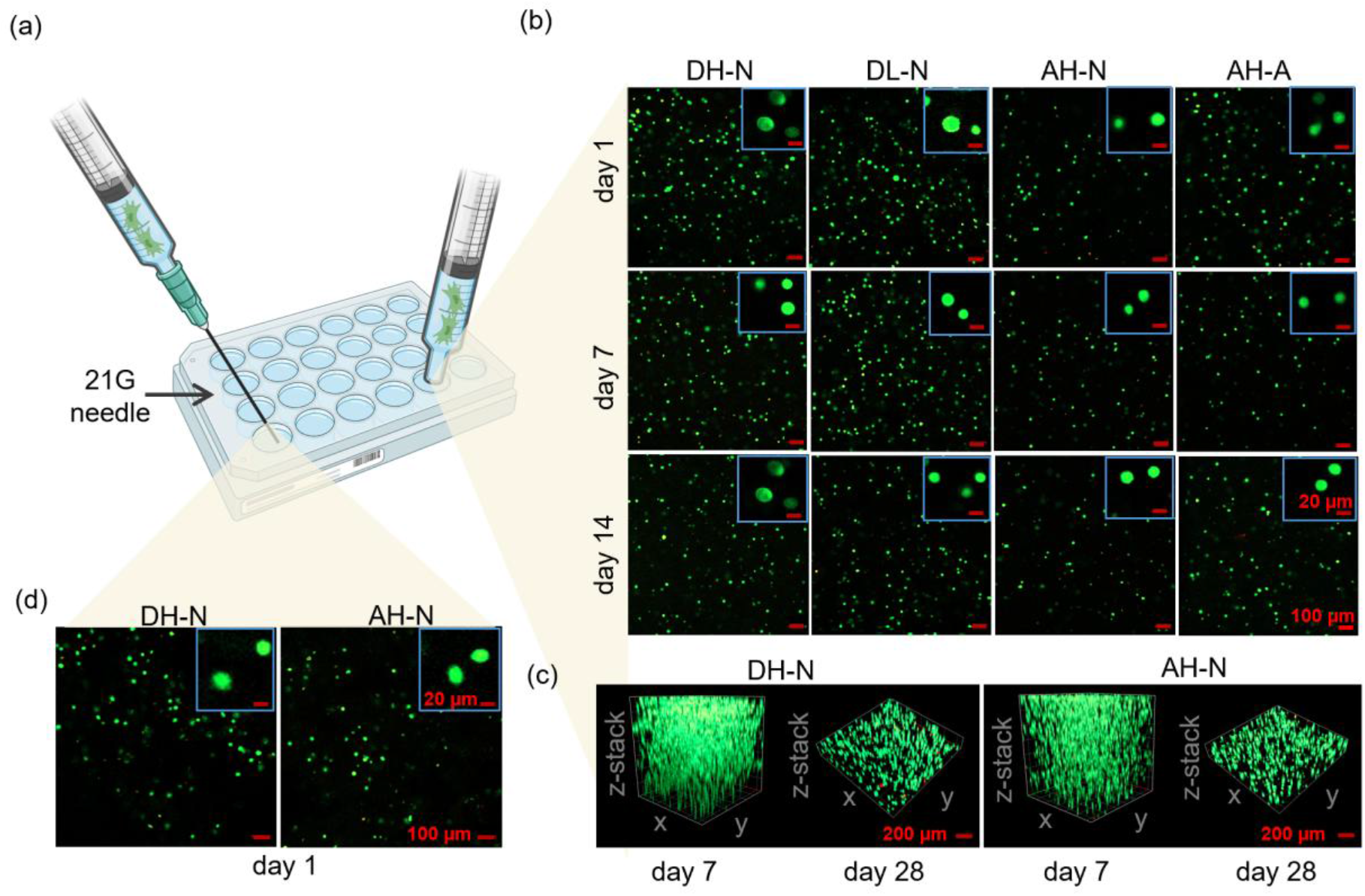
Viability and spatial distribution of MSCs-laden within acylhydrazone hydrogels. (a) Schematic illustration of direct casting of MSCs-laden hydrogels into a culture plate and of syringe injection through a 21 G needle prior to culture. (b) Confocal LIVE/DEAD images of MSCs-laden hydrogels after 1, 7, and 14 days of culture. Insets show representative higher-magnification views of encapsulated cells. (c) Three-dimensional confocal z-stack reconstructions of MSCs-laden DH-N and AH-N hydrogels after 7 and 28 days of culture. (d) Confocal LIVE/DEAD images of MSCs-laden hydrogels after syringe extrusion through a 21 G needle. Live and dead cells are shown in green and red, respectively. Scale bars: 100 µm in the main images and 20 µm in the insets b and d, and 200 µm in the 3D reconstructions c.

Confocal fluorescence imaging of MSCs-laden hydrogels transferred by direct casting to well plates (DH-N, DL-N, AH-N, and AH-A shown in Figure 11b and AL-A and AH-N in Figure S8, Supplementary Information) demonstrated high cell viability in all hydrogels throughout the culture period. In all formulations, the majority of laden cells exhibited green fluorescence at days 1, 7, and 14, indicating sustained cell survival after encapsulation and during prolonged culture, whereas only a limited number of dead cells were detected by red fluorescence. Interestingly, MSCs remained highly viable in AH-A hydrogels despite the mildly acidic pH of the OxA-H solution and AH-A hydrogel. This may be attributed to the short duration of cell exposure to the acidic environment (around 5 mins) after immediate immersion in culture medium, allowing rapid neutralization of the hydrogel pH.

Figure 11b and Figure S8 (Supplementary Information) show that all hydrogels supported a homogeneous distribution of MSCs throughout the hydrogel network without visible aggregation. In addition, Figure 11c shows 3D confocal reconstructions that further enable qualitative assessment of the spatial distribution of MSCs within representative hydrogels (AH-N and DH-N) on day 7 and day 28. In both systems, MSCs remained uniformly distributed throughout the hydrogel constructs for up to 28 days, with no evidence of sedimentation and clustering. Importantly, both AH-N and DH-N hydrogels supported a homogeneous distribution of MSCs throughout the hydrogel network without visible aggregation despite their distinct gelation kinetics (Figure 2c). This finding is noteworthy because rapid gelation of DH-N may restrict complete cell dispersion during mixing, whereas slow gelation of AH-N may allow cell sedimentation before sufficient network formation. The homogeneous distribution observed in AH-N suggests that the initial stages of acylhydrazone crosslinking were sufficient to immobilize cells and prevent sedimentation, even though the hydrogel required additional time to reach its network equilibrium. Equally, the homogeneous distribution in DH-N demonstrates that the encapsulation protocol provided effective cell dispersion within the Alg-ADH solution and sufficient time for homogeneous mixing with the OxD solution.

Injectable hydrogels need to protect cells from mechanical stress generated during the syringe-based administration. Therefore, the influence of injection on MSCs viability was evaluated by extruding cell-laden AH-N and DH-N hydrogels through a 21 G needle, followed by LIVE/DEAD staining 24 h post-injection. Figure 11d shows that the post-injection viability did not differ to that shown for directly casted hydrogels. The cells remained homogeneously distributed throughout the hydrogel network, with no apparent qualitative differences between the slowly gelling AH-N and rapidly gelling DH-N systems. The preservation of MSCs viability after syringe extrusion is consistent with previous reports on dynamic acylhydrazone hydrogels. Morgan et al. demonstrated that OxA-based dynamic hydrogels containing mixed hydrazone and oxime crosslinks could be extruded through a 24G needle while maintaining >99% viability of encapsulated human dermal fibroblasts within the hydrogel and >85% overall cell viability after injection.^[6b]^ Similarly, Leo et al. reported that a 1.5 wt% hyaluronic acid hydrogel crosslinked through hydrazone bonds exhibited shear-thinning and self-healing behavior, allowing extrusion-based 3D bioprinting while maintaining >80% viability of encapsulated fibroblasts post-printing.^[10a]^ Notably, the 1.5 wt% acylhydrazone-crosslinked hyaluronic acid hydrogel reported by Leo et al. required a low extrusion force, remaining below 10 N when extruded through fine 18, 23, and 25 G needles, similar to the present study. Thus, the combination of shear-thinning behavior (Figure 9c), rapid network recovery (Figures 5 and 9a), and syringe extrusion through a 21 G needle at a low extrusion force (Figure 9b) protected encapsulated MSCs from injection-induced mechanical damage.

Representative high-magnification images of MSCs shape are shown in the upper-right insets of Figure 11a. MSCs retained a predominantly rounded cell shape on days 1, 7, and 14, with no apparent differences between the fast- and slow-gelling hydrogel formulations. Likewise, no changes in MSCs shape were observed following syringe injection through a 21 G needle, as shown in Figure 11d. Previous study has demonstrated that MSCs shape in dynamic hydrazone hydrogels is regulated by matrix viscoelasticity and stress relaxation.^[8a]^ Specifically, hydrazone hydrogels formed using side-chain reaction of aldehyde- and hydrazide-functionalized polymers supported a predominantly rounded MSCs shape, whereas hydrogels based on star-shaped crosslinkers promoted a more elongated cell shape. Consistent with these observations, MSCs encapsulated within the side-chain acylhydrazone hydrogels investigated in the present study also maintained a predominantly rounded shape.

#### 3.11.2. Chondrocytes-laden hydrogels

The viability and spatial distribution of chondrocytes-laden hydrogels were evaluated using LIVE/DEAD staining and confocal microscopy. AH-N and DH-N hydrogel precursors were cast directly into well plates. Chondrocytes exhibited high viability and a homogeneous distribution throughout hydrogels volume, confirming the cytocompatibility of the hydrogels toward human chondrocytes (Figure S9, Supplementary Information), identically as observed for MSCs (Section 3.11.1). However, distinct differences in early chondrocyte shape were observed between the two hydrogels (Figure 12a), which is not observed for MSCs-laden hydrogels (Figure 11b). Figure 12a demonstrates that chondrocytes within DH-N displayed an elongated shape during the initial time following encapsulation, whereas chondrocytes in AH-N predominantly retained their characteristic rounded shape on day 1. Interestingly, after 14 days of culture, chondrocytes in both hydrogels exhibited a rounded shape. To capture chondrocyte shape from the point of encapsulation inside hydrogels, time-lapse quantitative phase imaging was performed, enabling continuous observation of individual cells during the first 18 h of culture using a label-free approach. The label-free approach enabled immediate visualization of native cell shape while avoiding staining and repeated washing procedures, thereby minimizing potential effects on early cell shape. Figure 12b shows that during the first 18 h following encapsulation, chondrocytes in AH-N are predominantly rounded in shape (Figure 12b, time points of 00:00, 10:00 and 18:00 h; Video S5-6), whereas chondrocytes in DH-N are elongated (Figure 12b, time points of 00:00, 10:00 and 18:00; Video S7-8). Hence, within the first 18 hours, there is no shape relaxation of chondrocytes immobilized in fast-gelling DH-N hydrogels. This early period is critical for establishing initial cell–hydrogel interactions, as chondrocytes begin depositing pericellular extracellular matrix within the first day after encapsulation.^[50]^

**Figure 12.**
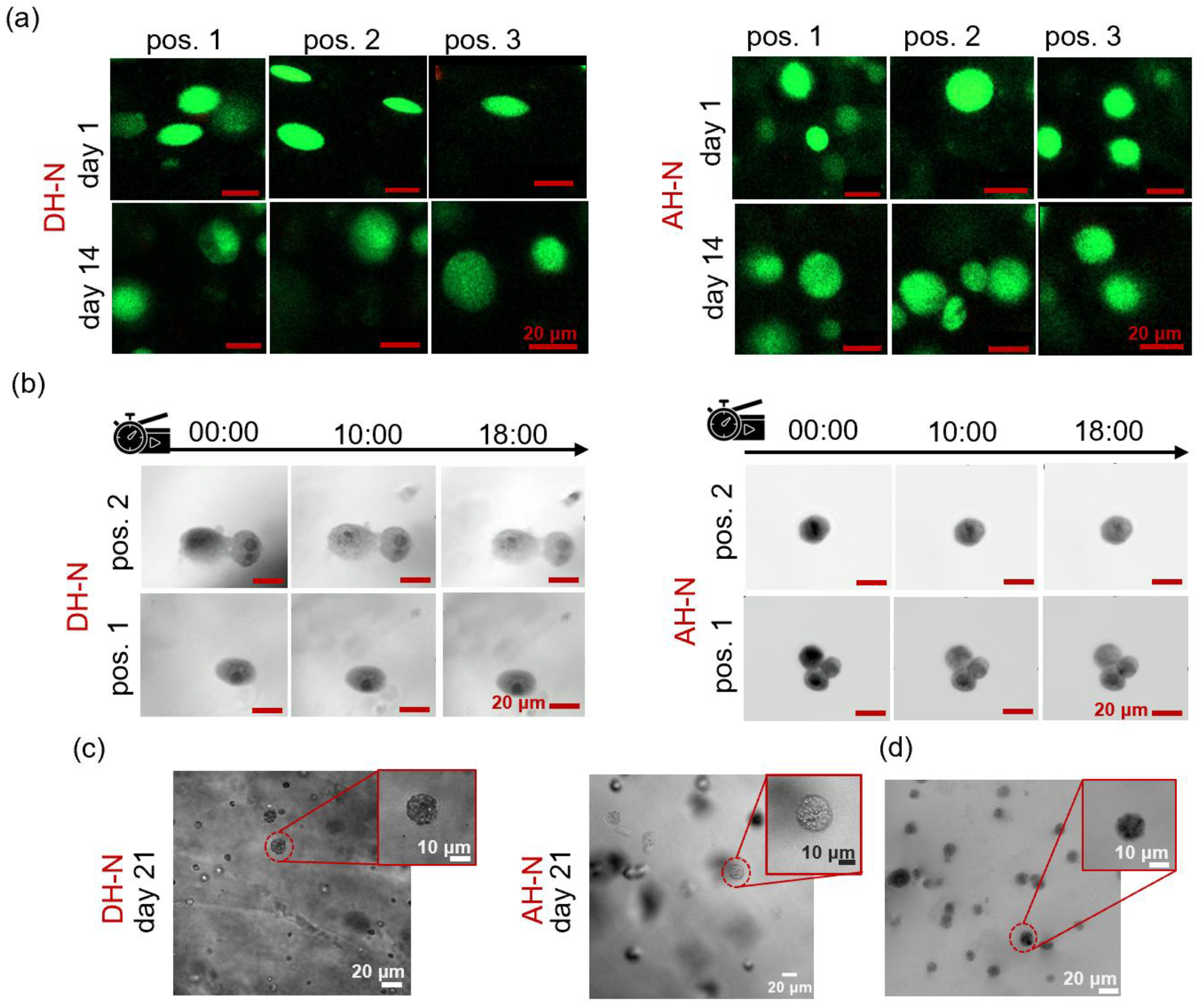
Viability and shape of chondrocytes-laden within acylhydrazone hydrogels. (a) Confocal fluorescence images of LIVE/DEAD-stained chondrocytes within hydrogels at day 1 and day 14, acquired at three different positions (pos. 1–3). Live cells are shown in green. (b) Time-lapse quantitative phase imaging showing early chondrocyte shape in DH-N and AH-N hydrogels at two representative positions (pos. 1 and pos. 2) over ∼18 h and (c) bright field image 21 days post-encapsulation. (d) Quantitative phase imaging of AH-N at day 21.

The chondrocyte deformation index (X/Y), calculated as the ratio of the minor (X) to the major (Y) cell axis, was used to quantify cell shape. A deformation index of 1 indicates a rounded cell, whereas lower values indicate increasing cell elongation. Confocal fluorescence imaging showed that chondrocytes encapsulated in AH-N maintained a nearly spherical shape throughout culture, with deformation indices of 0.98 ± 0.01 on day 1 and 0.99 ± 0.001 on day 14. In contrast, chondrocytes in DH-N exhibited marked elongation, with a deformation index of 0.44 ± 0.14 on day 1, and subsequently recovered to a nearly rounded morphology, reaching 0.95 ± 0.03 by day 14. Quantitative phase imaging showed a similar trend. In AH-N, the deformation index remained close to unity throughout the culture period, with values of 0.96 ± 0.04 on day 0, 0.96 ± 0.04 after 18 h, and 0.95 ± 0.07 on day 21. In contrast, chondrocytes in DH-N displayed a lower deformation index immediately after encapsulation (0.72 ± 0.08 on day 0), which persisted during the first 18 h, before recovering to a rounded shape by day 21 (1.00 ± 0.01). These findings are consistent across both fluorescence and quantitative phase imaging, demonstrating that chondrocytes initially undergo transient elongation in DH-N but progressively regain their native rounded shape during culture.

AH-N and DH-N hydrogels exhibited comparable stiffness (Figure 4b and Figure 6b), viscoelastic properties (Figure 4a and c), and stress relaxation behavior (Figure 7). Hence, the substantial differences in gelation kinetics are seen responsible for the distinct early cell shape in these hydrogels. AH-N of slow gelation (Figure 2c) provides a gradual transition from the liquid precursor state to the fully crosslinked hydrogel. A slow crosslinking kinetics reduces mechanical confinement during encapsulation and mixing, allowing cells to adapt progressively to the evolving network environment and, thereby, preserving the rounded shape of chondrocytes. In contrast, DH-N exhibits rapid gelation within approximately 2 min (Figure 2c). This near-instantaneous network formation likely exposes encapsulated cells to higher transient mechanical stresses during encapsulation, which induce their elongation in the early stages after encapsulation. Unlike MSCs, which maintained a rounded morphology irrespective of gelation kinetics (Figure 11b and Figure 11c), chondrocytes exhibited transient differences in morphology during the early stages of hydrogel formation, suggesting that the influence of gelation kinetics is cell type-dependent. Chondrocytes encapsulated in AH-N maintained a rounded cell shape throughout the culture period (Figure 12). Importantly, chondrocytes encapsulated in DH-N recovered a rounded shape during culture (Figure 12c). This recovery suggests that the initial chondrocyte deformation was reversible and that the dynamic hydrogel network enabled adaptation of cell shape over time. A previous study demonstrated that dynamic alkylhydrazone-based hydrogels facilitated recovery of the native rounded chondrocyte shape during compressive deformation, whereas elastic benzylhydrazone-based hydrogels significantly limited morphological recovery.^[7]^ However, in that study, cell deformation was induced by externally applied mechanical loading, and shape recovery occurred rapidly after removal of the applied stimulus, regardless of hydrogel viscoelasticity. In contrast, no external mechanical deformation was applied in the present study. A round shape of chondrocytes during culture is characteristic for chondrocyte phenotype, which was also reported for 2D culture.^[46]^ Our findings suggest that dynamic viscoelastic hydrogels support the recovery of chondrocyte shape following deformation associated with hydrogel formation.

## Conclusions

A major challenge in the development of injectable dynamic biomaterials is the lack of predictive understanding of how polymer structure and reaction conditions govern gelation kinetics and, ultimately, how these kinetic processes translate into mechanical performance and biological function. This work addresses this challenge by investigating acylhydrazone-crosslinked hydrogels formed from adipohydrazide-functionalized alginate (Alg-ADH) and oxidized polysaccharides with distinct backbone chemistries, namely oxidized alginate (OxA) and oxidized dextran (OxD).

By integrating variations in polysaccharide structure, degree of oxidation, and pH, we establish a quantitative structure–kinetics–property–function framework that identifies gelation kinetics as the critical mechanistic link between molecular design and hydrogel performance. This kinetic perspective represents a key advance of the study, revealing that oxidized polysaccharide backbone chemistry is the primary determinant of network formation. Specifically, OxD-based hydrogels undergo rapid, largely pH-independent gelation, whereas OxA-based systems exhibit pronounced pH-sensitive and substantially slower network formation under physiological pH, ultimately governing their mechanical stability and cellular responses.

These kinetic differences propagate across hierarchical levels of material behavior. Whereas the gelation behavior of OxD-based hydrogels remained unaffected by pH, OxA-based hydrogels exhibited not only accelerated gelation but also increased stiffness and enhanced stress relaxation, suggesting that pH influences both the kinetics and the equilibrium of the OxA-based acylhydrazone network rather than acting solely as a kinetic catalyst. In contrast, at neutral pH, OxA- and OxD-based hydrogels displayed markedly different gelation kinetics, while converging to comparable stiffness and stress-relaxation behavior, suggesting that the OxD backbone primarily facilitates rapid network formation without substantially altering the final mechanical response. Together, these findings demonstrate that both backbone chemistry and pH regulate acylhydrazone network formation, with backbone chemistry primarily controlling gelation kinetics and pH modulating both the kinetics and equilibrium mechanical properties of the resulting OxA-based hydrogels. Importantly, all formulations retain essential functional attributes, including shear-thinning injectability, low extrusion forces, and rapid self-healing with efficient recovery after mechanical deformation.

Long-term culture studies further demonstrate that backbone chemistry governs structural stability and mechanical properties, with OxD hydrogels maintaining stability and showing reinforcement under dynamic loading, whereas OxA networks exhibit faster degradation under physiological environment. Biological assessments confirm high cytocompatibility across all formulations; however, gelation kinetics critically regulate early chondrocyte morphology, with slowly forming OxA hydrogels preserving a rounded phenotype and rapidly gelling OxD networks inducing transient cell elongation, while mesenchymal stem cells remain largely unaffected.

Overall, this study establishes gelation kinetics as a central design parameter linking polymer structure to function in dynamic covalent hydrogels. The resulting structure– kinetics–property–function framework provides a rational basis for engineering injectable, mechanically adaptive biomaterials and offers important design principles for their application in cartilage tissue engineering and regenerative medicine.

## Acknowledgments

We acknowledge the Biophotonics Core Facility, CEITEC Brno University of Technology, Czech Republic supported by MEYS CR (LM2023050 Czech-BioImaging). AH and JZ acknowledge the COST Action CA21110─Building an Open European Network on Osteoarthritis Research (NetwOArk) funded by the European Union and the European Commission under the European Cooperation in Science and Technology Programme (COST). FKA acknowledges the Presidium of the Slovak Academy of Sciences for partially supporting this work through the Doktogrant program (Project APP0667).

## Data availability

Supporting data associated with this article will be made publicly available in a Zenodo repository upon acceptance. The corresponding DOI will be provided during the proof-reading stage.

